# Spatial frequency tuning of perceptual learning and transfer in global motion

**DOI:** 10.1101/372458

**Authors:** Jordi M Asher, Vincenzo Romei, Paul B Hibbard

## Abstract

Perceptual learning is typically highly specific to the stimuli and task used during training. However, recently it has been shown that training on global motion can transfer to untrained tasks, reflecting the generalising properties of mechanisms at this level of processing. We investigated a) if feedback was required for learning when using an equivalent noise global motion coherence task, and b) the transfer across spatial frequency of training on a global motion coherence task, and the transfer of this training to a measure of contrast sensitivity. For our first experiment two groups, with and without feedback, trained for ten days on a broadband global motion coherence task. Results indicated that feedback was a requirement for learning. For the second experiment training consisted of five days of direction discrimination on one of three global motion tasks (broadband, low or high frequency random-dot Gabors), with trial-by-trial auditory feedback. A pre- and post-training assessment was also conducted, consisting of all three types of global motion stimuli (without feedback) and high and low spatial frequency contrast sensitivity. We predicted that if learning and transfer is cortically localised, then transfer would show specificity to the area processing the task (global motion). In this case, we would predict a broad transfer between spatial frequency conditions of global motion only. However, if transfer occurred as a result of backward generalisation, a more selective transfer would occur matching the low-pass broadband tuning of the area processing global motion. Our training paradigm was successful at eliciting improvement in the trained tasks over the five days. However, post-training transfer to trained or untrained tasks was only reported for the low spatial frequency trained group. This group exhibited increased sensitivity to low spatial frequency contrast, and an improvement for the broadband global motion condition. Our findings suggest that the feedback projections from global to local stages of processing play a role in transfer.

## Introduction

Perceptual learning has attracted much attention as a potential tool to aid recovery of lost visual function for clinical populations [1]. The success of perceptual training in amblyopia [2–4], presbyopia [4] and cortical damage [5–7] has demonstrated sensory plasticity in adulthood. This evidence contradicts the position that sensory development is restricted to a critical period early in life [8, 9] and that the visual system is hard-wired in mature systems [10]. While it has repeatedly been established that training can improve perceptual abilities [11], these benefits tend to be highly specific for both the perceptual features of the stimuli [12–14] and the behavioural task used in training [15]. This specificity severely limits the effectiveness of perceptual learning as a general therapeutic tool. Resolving the conditions under which learning is tied to the features and tasks used in training, and how much it can generalise to new tasks and stimuli, is imperative for understanding the mechanisms of perceptual learning [11, 16]. Our study aims to identify the relative location of one of the mechanisms involved when learning direction discrimination for a global motion coherence task. We do this by evaluating the spatial frequency tuning of improvements in performance for trained and untrained tasks. For the purpose of this paper we dissociate between models and mechanisms of perceptual learning using two general categories i) the How and ii) the What and Where. Models of perceptual learning aim to resolve ‘how’ learning occurs. Understanding which mechanisms underlie this learning address the ‘what’ and ‘where’ (and possibly ‘why’) questions. In this way we consider a mechanism a feature or component of the learning model [17].

### Models of Perceptual Learning - How

The ‘how’ is often the primary focus for research and there are, broadly speaking, two predominant models of perceptual learning. The first explains perceptual learning in terms of a change in the neurons that code for that feature [18, 19]. The second position argues that learning is a result of the change in weights of readout between the sensory representation and the decision units [20–24]. The first position developed as a result of accumulating evidence that improvements were highly specific for the features of the stimuli used for training. Early studies identified that learning was specific for orientation [13, 25, 26], spatial frequency [11, 13, 27], direction of motion [15], retinal location [14, 24, 26, 28–30] of stimuli and the eye to which they are presented [14, 24, 29]. On the basis of this feature-specificity it has been argued that the underlying brain area responsible for the learning process must be within the primary visual cortex [14] where the receptive fields of cells display a high degree of specificity. The implication of this position is a potential degree of plasticity in individual neurons within an area previously thought to be incapable of structural change [14, 19, 31]. Although the failure of perceptual learning to transfer across retinal location, orientation and other stimulus dimensions implicates V1, learning is far more complex than can be explained by the simplicity of the stimulus driven, bottom-up, visual representation model [32]. A model of learning also needs to include top-down factors such as attention (see [31] for a review), confidence in decision making [33, 34] and knowledge and understanding of the task [35]. The second position proposes that repeated experience alters the connections between lower visual areas and the high level areas where the perceptual decision is then made [36]. Rather than a physical change in the sensory neuron, learning represents a change in the weights of connections between the input, sensory encoding layer and the classification layer at which the decision is made [20–24].

One consistent aspect across models, is that perceptual learning is likely to occur as a result of exploiting correlations between neural responses [19].

Behavioural studies of perceptual learning are unable to distinguish between learning as a result of changes in sensory encoding, or as a result of a reweighting of the connections between sensory and decision-making stages of processing. However, learning effects on the structure of receptive fields of visual neurons have been measured using single cell recordings. Changes in orientation and spatial frequency tuning in V1 [37, 38] and V4 [39], and increased sensitivity to motion coherence in V5 [40], have all been found as a result of perceptual learning. In addition, neuroimaging in human observers has indicated a reorganisation of the visual cortex after training [41–43]. However, this in itself cannot dissociate between learning models [32, 35, 44]. Thus, it is generally judged that physiological models are not sufficient to explain the improvements in task performance exclusively [21, 45].

### Mechanisms of Perceptual Learning - What, Where, Why

In order to begin to unravel the ‘what’, ‘where’ and ‘why’ questions, the behaviours of the system in question need to be understood, and the sensory output mapped to the behavioural output [17]. In this section we will consider some of the behaviours associated with the learning process.

#### Specificity

A hallmark behaviour of perceptual learning is its specificity to the task, and features, of training. Early psychophysical research investigated learning and transfer using lower-level visual tasks such as texture discrimination [14] and motion discrimination [46] that match the receptive fields of neurons in the lower visual processing areas. At the lowest level, V1 and V2 represent simple visual dimensions, such as the position, orientation and scale of local image features [12]. The receptive fields of cells in V1 are small and only respond to a very restricted area of the visual field [47–49]. Furthermore, V1 neurons are sharply tuned to orientation [50, 51], spatial frequency [50, 52] and direction [53, 54]. For example contrast detection, which is a low-level visual task, is proposed to depend on processing in V1. The pathway for contrast detection and discrimination is believed to develop during childhood and adolescence, reaching its peak in adulthood, followed by a decline in late adulthood [55]. Early psychophysical studies found little perceptual learning for contrast-dependent stimuli [18]. More recently, learning has been found to occur as long as ‘enough training’ is provided [56]. The improvement in contrast sensitivity has been found to be highly tuned to spatial frequency, specific for the trained eye and retinal location of training, but not selective for orientation [27, 56–61]. Conversely, the receptive fields in higher cortical areas, such as those found in V4 and V5, are larger than those of V1 and are less dependent on location, retinal size, viewpoint, and lighting [47, 52, 62–66].

#### Transfer

More recently, a number of studies have demonstrated transfer following training using a double-training protocol [59, 61, 67]. Double-training involves overt training on a task-relevant feature in addition to exposure to a task-irrelevant feature [68]. Contrast sensitivity, historically specific to the retinal location of of training was found to transfer between retinal locations when observers were trained on two simultaneous tasks [59]. Observers were required to detect a relevant feature (contrast) at one retinal location, followed by detection of an irrelevant feature (orientation) at another location, and a full transfer of contrast sensitivity to the untrained location was found [59]. Similarly, observers who were trained to discriminate orientation and simultaneously exposed to a second orientation experienced a full transfer of improvement in orientation sensitivity as long as training preceded exposure, rather than being simultaneously presented [61]. Zhang et al. (2010) [61] proposed a rule-based learning model to account for the transfer, where higher level decision units learn and re-weight the V1 inputs. They proposed that the absence of functional connections to the untrained orientation or location prevents any potential reweighting. Double training, through exposure, activates the functional connections at the new locations or orientations, enabling transfer. This suggests that there may not be a straightforward correspondence between stimulus features and the neural loci involved in performing the task. Double-training has been a successful paradigm in obtaining evidence of transfer [59, 69, 70], however task difficulty during training, and observer confidence still play a vital role in the learning and transfer process [34, 71]. An early theoretical model of transfer is the Reverse Hierarchy Theory [72]. This model proposes that transfer is a top-down process, and the degree of transfer is dependent on the characteristics of the receptive fields involved in performing the training task. Transfer occurs as a result of modification of neurons found in the higher cortical levels, where receptive fields generalise. In contrast, specificity, occurs at the lower cortical levels where the receptive fields are localised [72]. As established, learning is highly specific to the features of the stimulus and reflects the tuning properties of the relevant receptive fields. However, what specificity is expected for tasks that are not processed in early visual areas? For example, perceptual learning improves detection and discrimination in tasks using global form [73] and global motion [46, 74, 75]. These kinds of tasks are known to be processed at higher levels of the visual cortex, for example areas V3 [76], V4 [73] and V5 [77, 78]. The Reverse Hierarchy Theory predicts that the feedback connections from higher levels back to early visual processing areas may be involved in facilitating learning that transfers, and that the key to understanding specificity and transfer lies in the hierarchy of processing and the feedforward and feedback connections between them [33, 72]. Based on this, tasks that require higher level integration or segregation (global processing) may therefore play an important role in creating learning that generalises to other tasks and stimuli [33].

#### Local vs Global Processing

The receptive fields of neurons in higher cortical areas integrate information to represent global stimulus properties [19, 79–82]. Global processes are investigated using stimuli or tasks that can only be resolved through integration and segregation of coherent or conflicting information [81–83]. Cells higher in the processing cortical hierarchy are involved in the perception of global aspects of an image and generalise across individual features such as spatial frequency and location. Based on the predictions of the Reverse Hierarchy Theory, modification of these generalising receptive fields may produce perceptual learning that also generalises over these stimulus parameters. Since perceptual learning occurs for both global motion [46, 75] and form tasks [83, 84], this suggests that learning is not restricted to the initial encoding of information in V1, and can occur at higher levels of cortical processing. Learning that involves higher level global aspects of perception is of particular interest since it has the potential to produce more generalisable improvements [33]. Huxlin et al. (2009) [5] suggested that extensive training using global motion may result in general improvements in visual sensitivity. They postulate that the improvements may be a result of the feedback connections from higher brain areas back to V1, where training may stimulate and reactivate “intact islands” of activity.

### Testing models of perceptual learning of global motion

Paradigms to evaluate the transfer of learning based on features of the task may shed light on the location of the learning mechanisms involved. In order to behaviourally disassociate the location of a learning mechanism, Petrov et al., [36, 44] suggest using perceptual learning experiments in which the sensory representations required for two tasks are the same, but the decision stages differ, or where both sensory representations and the decision stage are common to both. Training using a global motion task has been found to transfer between eyes [15, 46]; increase sensitivity for detection and discrimination [5]; help recover some of the blind field for cortically blind subjects [5]; and reduce contrast thresholds for drifting stimuli [75]. Transfer of learning has also been reported in a control population [75], where a post-training improvement was found for contrast sensitivity. This is a particularly interesting result when taking into account that a globally processed trained task improved contrast detection, which is known to be processed in an early visual locations such as 4C*α*, where cells are tuned for spatial frequency but not orientation [27]. The success in obtaining improvements in untrained tasks/ features makes global motion an interesting task to investigate this further.

#### Global Motion Processing

The perception of motion is hypothesised to occur as a two stage process [80, 85]. At the first stage, spatial frequency- and orientation-tuned mechanisms in V1 encode the motion signals that occur locally within the receptive fields of individual neurons [54, 81]. However, local and global stages of motion processing differ in their spatial frequency tuning. In order to process more complex motion, ambiguous or conflicting signals from the first stage need to be integrated over a wider spatial area to provide a global representation of motion and velocity [86, 87].

A number of areas within the visual cortical hierarchy play a functional role in processing motion. Areas V2 and V4 have a role in processing moving orientation signals [88, 89]. V3A also plays a role in several aspects of motion processing [62], with 76% of neurons being selective for orientation and 40% showing strong direction selectivity. However, evidence from lesion studies [90, 91], extra-cellular recordings [77, 92] and neuroimaging in humans [93] support area V5 as a brain area that is heavily involved in processing global motion [77, 85, 90–92, 94]. Most neurons in V5 are strongly direction selective [93, 95], and the evidence for the role it plays in spatially integrating motion signals is well supported by non-human primate data, and neuroimaging studies in humans [47, 77, 96, 97]. Receptive fields in V5 can be up to tenfold larger than those in V1 [65], with broad spatial frequency and orientation tuning, allowing them to sum the responses of V1 neurons, across space, orientation and spatial and temporal frequency [49].

#### Spatial frequency tuning of global motion processing

Psychophysical studies have shown that global motion detectors have relatively broad spatial frequency tuning [80]. This is consistent with single-cell recordings from area V5 in marmoset monkeys that exhibit bandpass spatial and temporal frequency tuning, with a preference for low spatial frequencies [96]. Amano et al. (2009) [79] suggested that, within V5, there is a “motion-pooling mechanism” with broadband, low-pass tuning. Pooling of visual sensory information has also been found in studies investigating transfer in binocular stereopsis [98]. The neural mechanisms of stereoscopic vision are also known to be processed on multiple levels of the visual hierarchy [99], and the pooling of information across spatial frequency mechanisms has been proposed as an important step in estimating depth [100–103]. In the same way, pooling across low frequencies is likely an important step for integration and segregation of coherent motion. Furthermore, a recent fMRI study [104] reported that areas V5+ and V3A both exhibit an attenuated response for high spatial frequencies.

#### Global motion and the reverse hierarchy in practice

The Reverse Hierarchy Theory predicts that an increase in sensitivity at lower cortical areas, as a result of the feedback connections from higher cortical areas, would be dependent on the frequency tuning of the global motion detectors [72]. Feedback connections are argued to be fundamental to efficient cortical organisation, and of all the feedback systems terminating in V1, the connections from V5 to V1 have been found to cover the most territory [105]. Furthermore, these feedback (or re-entrant) connections terminate in different combinations within various layers throughout V1 [65, 105] and are rapidly updated [106, 107]. Recordings from macaque monkeys have suggested that there is almost no delay for information processed in V5 to be fed back to lower areas, and it has been proposed that the feedback from V5 is present prior to the bottom up information from the feedforward connections [106, 107]. This has recently been supported by evidence of an independent motion pathway in humans, finding a direct link from lateral geniculate nucleus (LGN) to V5 [108]. Projections are targeted retinotopically to V1 in locations lying within the V5 receptive field [65]. Thus, as V5 responses are influenced by the bottom-up responses from V1, the top-down responses from V5 shape the responses of the V1 cells which provide the sensory input [65]. Sillito et al. (2006) [65] suggest that the “iterative interaction” between the two stages could account for the selectivity at both levels. Backward projections are a theoretically plausible route for perceptual learning, and are consistent with the predictions from the Reverse Hierarchy Theory. Overall this suggests a theoretically important role for top-down information in obtaining transfer from global motion [105].

Transcranial Magnetic Stimulation (TMS) has been used to stimulate the re-entrant connections from V5 to V1, enhancing the perception of global motion [109]. Using a novel paired cortico-cortical TMS protocol (ccPAS) to induce Hebbian plasticity, observers’ thresholds for motion detection were reduced when the feedback connections from V5 to V1 were stimulated. However, the improvement in perception was critically dependent on the timing and direction of stimulation. There was no change when the feedforward connections from V1 to V5 were stimulated [110]. This also suggests that these re-entrant connections are malleable [110]. Using a similar method, a direction-selective improvement was induced by pairing subthreshold stimulation with the simultaneous presentation of direction-specific moving stimuli [111]. This provides additional support for the accumulating evidence that the re-entrant connections from direction-tuned neurons in areas such as V5 play a role in perceptual learning in global motion coherence tasks.

### Our Study

Given the differences in spatial and temporal frequency tuning of processing between the local and global levels [50, 77, 96], measuring the tuning of training for these dimensions allows us to understand the role of each level in global motion learning.

Prior to collecting the data for our main study, we also questioned whether trial-by-trial feedback was a requirement for learning. Therefore, we first investigated the necessity of feedback for perceptual learning to occur for our specific stimuli.

#### Experiment 1

The specific nature of feedback and its role in perceptual learning is unclear, and external feedback has been shown to improve learning and increase efficiency [112, 113]. However, some studies have found that learning occurs without external feedback [11, 15, 30, 36, 78, 114]. Recently, it has been found that interleaving high accuracy (easy) trials and low accuracy (difficult) trials resulted in perceptual learning without the need for feedback, even on difficult trials [114]. Based on these results we predicted that we should find learning in both conditions, as long as easy and difficult trials were interleaved. As detailed in the following sections, our study found that learning did not occur in the condition where no feedback was provided, even with easy trials. Learning only occurred for the feedback condition. With this in mind our design for experiment 2 included feedback during training, but no feedback when testing.

#### Experiment 2 - Main Study

The purpose of this study was to extend the pre- and post-training measures captured by Levi et al. (2015) [75]. Thus, as well as testing for transfer to contrast sensitivity, we included measures of transfer to the trained and untrained global motion frequencies.

Transfer from global motion could involve a combination of reweighting that includes the global motion mechanisms in higher areas such as V5, and the re-entrant connections from global mechanisms to local mechanisms. Differences in the spatial frequency tuning at the local and global stage mean that these models make different predictions for the transfer of learning across spatial frequency, and different psychophysical tasks (Fig. 1).

**Fig 1.**
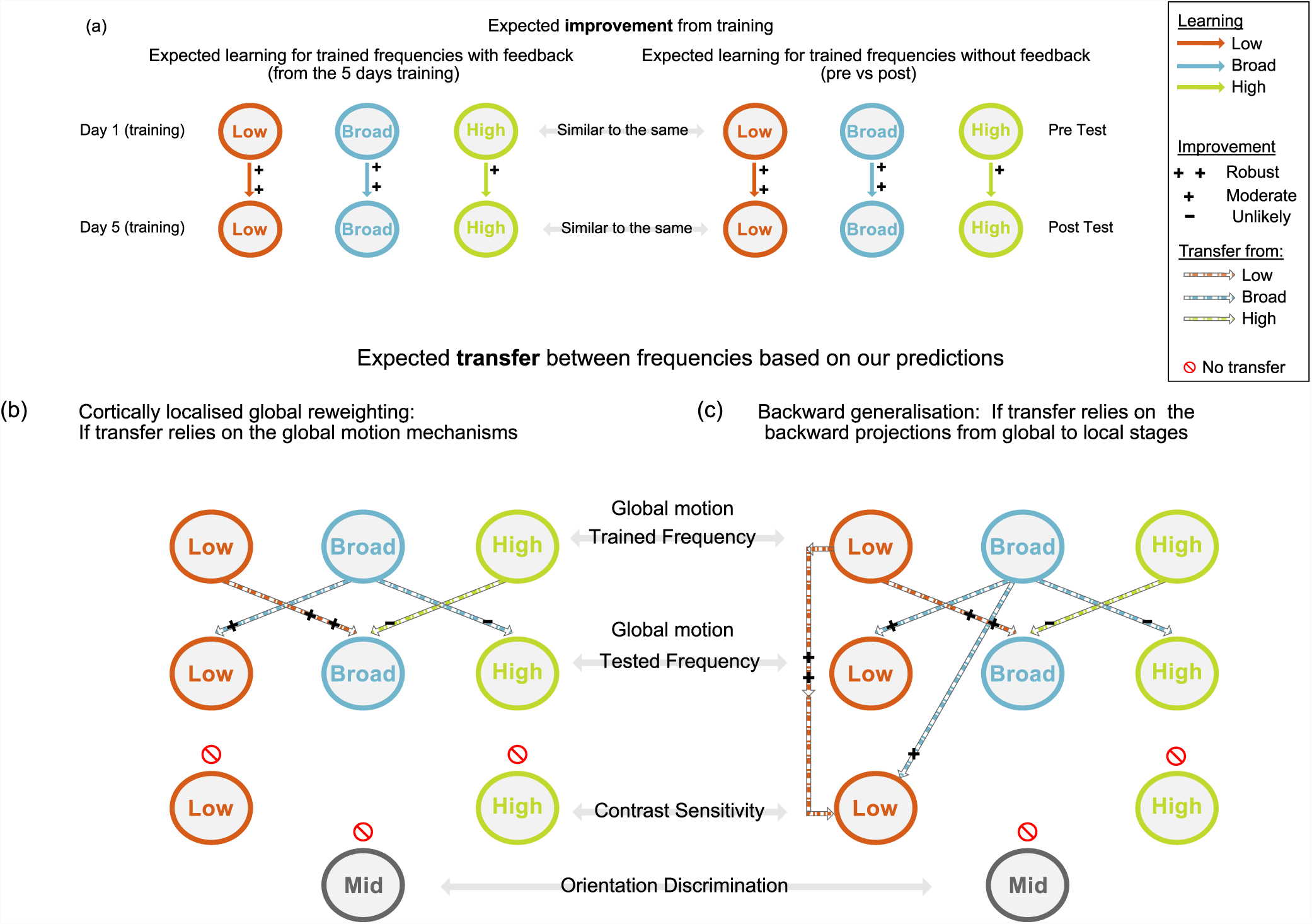
(a) **Expected improvement from training:** We would expect robust improvement on frequency specific motion stimuli over the 5 days training, and for this to be reflected at the post-assessment as an improvement compared to the pre-assessment results. Predictions of transfer (b) **Cortically localised global reweighting:** In this case we may predict that transfer from frequency specific global motion would reflect the broad, approximately low-pass spatial frequency tuning of global motion detectors in areas such as V5 and V3/V3A. There would thus be most transfer when both training and test stimuli contained low spatial frequency components (c)**Backward generalisation:** Should transfer occur as a result of the backward projections from V5 to V1 (as predicted by the Reverse Hierarchy Theory [33]) we would therefore also expect robust transfer from low to low frequency stimuli; modest transfer from low to broad, and broad to low and broad to broad, with an unlikely but possible transfer from high to broad, and broad to high. Importantly, should there be transfer to contrast sensitivity we would expect this to reflect the low-pass spatial frequency tuning of global areas.

### Predictions

#### Cortically localised global reweighting: If transfer relies on global motion mechanisms

In this case learning, and transfer of learning, for global motion would reflect the broad, approximately low-pass spatial frequency tuning of global motion detectors in areas such as V5 and V3/V3A. There would thus be most transfer when the training and test stimuli contain low spatial frequency components (robust transfer is predicted from low frequency to low frequency conditions; modest transfer from low to broad frequency stimuli, and from broad to low and broad stimuli, and an unlikely but possible transfer from broad to high and high to broad.). No transfer to contrast sensitivity would be predicted, since learning would result in changes in weightings between direction-tuned global motion mechanisms, and higher-level decision stages.

#### Backward generalisation: If transfer relies on feedback projections from global to local stages of processing

If transfer occurs broadly across tasks and levels in a top down manner, as predicted by the Reverse Hierarchy Theory [33], we would predict transfer to be restricted by the frequency tuning of the global motion detectors, but to transfer more broadly across tasks. We would expect most transfer when both training and test stimuli contained low spatial frequency components (robust transfer from low to low frequency stimuli; modest transfer from low to broad, and broad to low and broad to broad, with an unlikely but possible transfer from high to broad, and broad to high). We would also expect transfer to contrast sensitivity, and for this to show the same low-pass spatial frequency tuning. This would also be consistent with the transfer found by Levi et al. (2015) [75].

Finally, a mid frequency orientation discrimination condition was included as a control task. Since none of the motion training stimuli contain any orientation information, we predicted no transfer to occur from any trained condition or either level of processing.

## Methods and Materials

### Observers

24 observers for Experiment 1 and 30 (new) observers for Experiment 2 were randomly and evenly assigned into groups. For experiment 1 there was a feedback and a no-feedback group. For experiment 2 groups were categorised by the spatial frequency of training (broad, low, high). All observers were employees or students from the University of Essex and self declared as having normal or corrected-to-normal vision. All work was carried out in accordance with the Code of Ethics of the World Medical Association (Declaration of Helsinki). The study procedures were approved by the University of Essex Ethics Committee (JA1601). All observers gave informed written consent and were either paid or received course credit for their participation.

### Experiment 1

#### Apparatus

Stimuli were generated and presented with Matlab 2015a using the Psychophysics Toolbox extensions [115–117]. The broadband global motion stimuli were presented using a 27” 2.7 Ghz iMac running OSX 1.9.5 iMac with a display resolution of 2560 x 1440 pixels and a 60 Hz refresh rate. Viewing distance was 450mm, the stimuli subtended a visual angle of 66.8°, and one pixel subtended 1.77 arc minutes.

#### Stimuli and Procedure

Global motion stimuli contained 100 Gaussian elements, each with a standard deviation of 6.8 arc minutes (see Fig. 2a). Elements were presented within a mid-grey rectangle measuring 17.6° x 17.6° on a mid-grey background. Elements moved 5 pixels per frame, and each element moved a fixed distance of 8.8°. Dots wrapped around the rectangle when approaching the edges.

**Fig 2.**
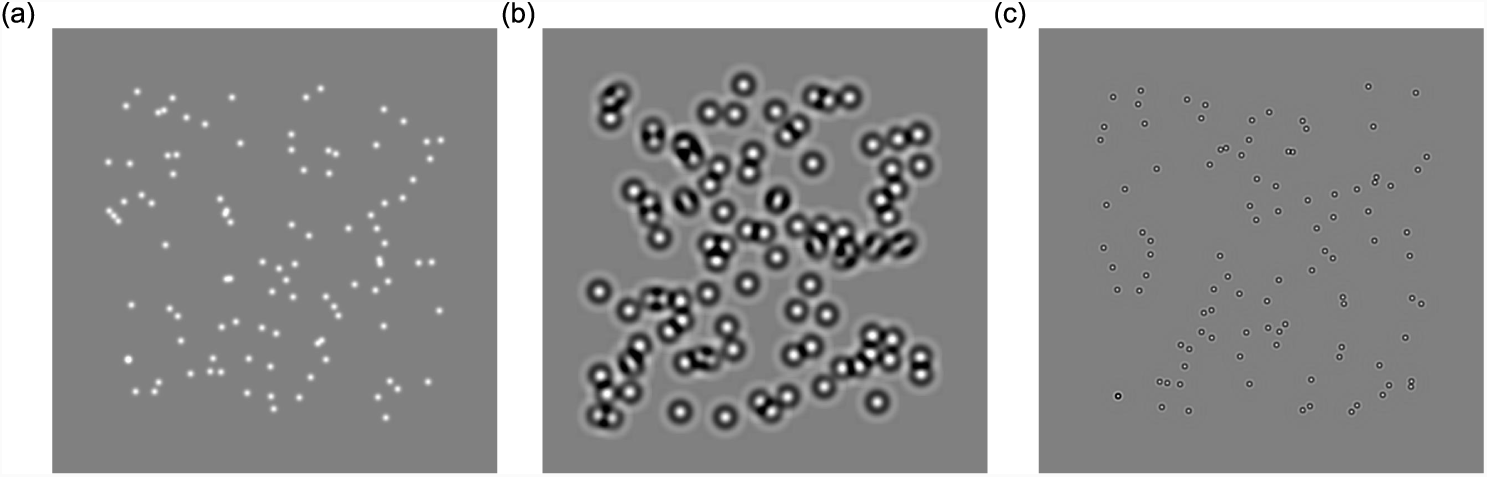
Global motion stimuli, at broad (a), low (b) and high spatial frequencies (c). For all stimuli the spatial frequency and speed of motion were held constant.

Global motion stimuli were based on the task designed by Williams and Sekuler (1984) [118]; dots moved around the screen at 7 levels of coherence (180°; 300°; 330°; 340°; 345°; 350°; 355°). Lower numbers depict higher coherence, and higher numbers depict lower coherence and thus more randomness. At each level of coherence each dot moved a fixed distance within the defined arc of degree of coherence, while moving in a leftwards or rightwards direction (see Fig. 3). All tasks were presented as two-alternative-forced-choice (2AFC) decisions (left or right response), using the method of constant stimuli (MOCS). Observers responded by pressing the left or right arrow key associated with perceived leftwards or rightwards motion. When feedback was present, this was provided as an immediate auditory beep after each trial, a high pitched tone for a correct response (2000Hz for 10ms), and a low pitched tone for an incorrect response (200Hz for 40ms). Each day 40 repetitions of each level of coherence was presented for a total of 280 trials.

**Fig 3.**
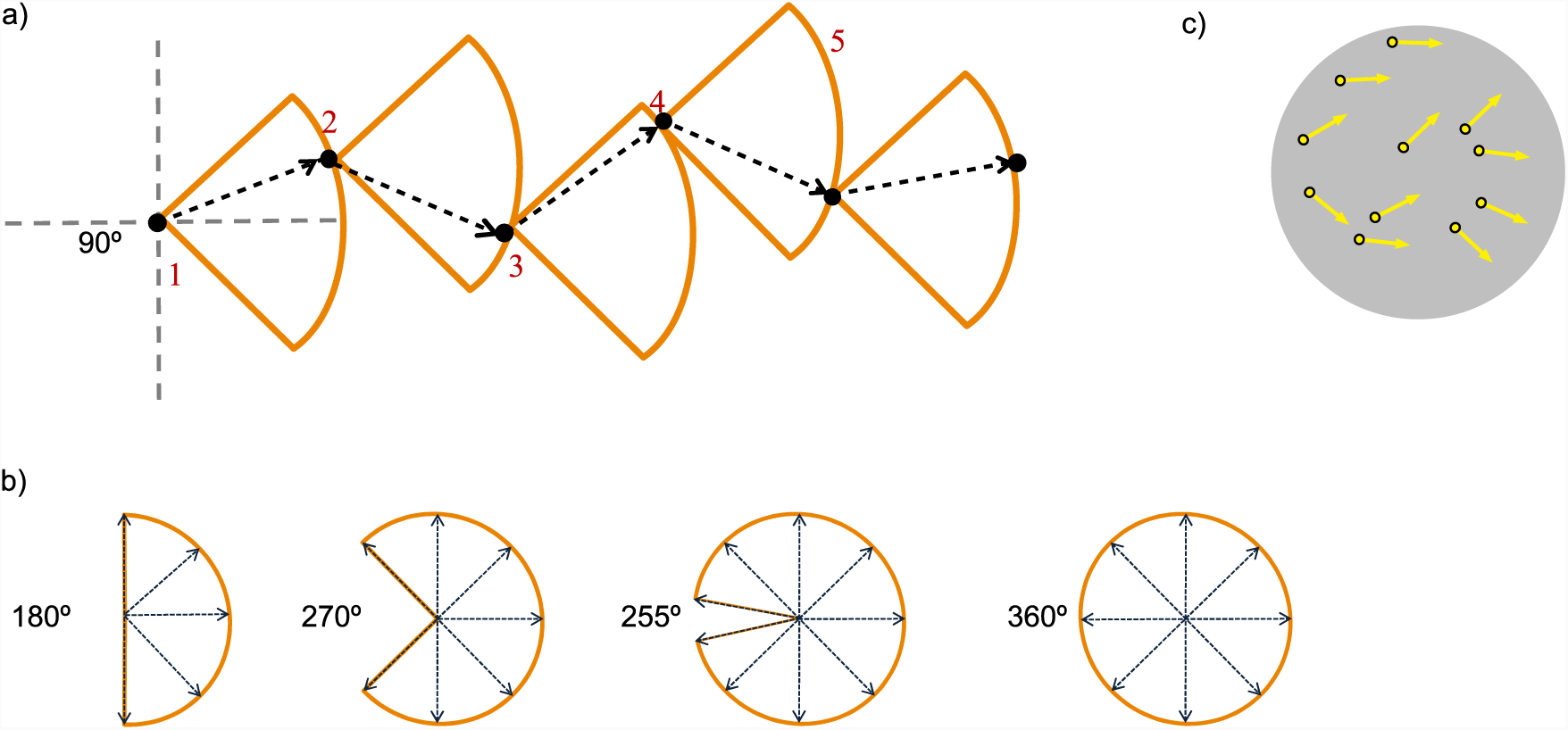
Equivalent noise global motion stimuli a) The random walk that creates the motion of dots with a 90° arc (for descriptive purposes only). The arc defines the potential area of movement each dot could take at that level. The higher the degree of arc, the lower the coherence. The arrow indicates the actual trajectory and motion of the dot on each step. b) Potential trajectories for the random walk for levels with 180°, 270° 255°and 360° arc. c) An schematic example of an equivalent noise global motion stimulus. All dots contribute equally to the signal and the noise (drawn from a distribution between 0° and 360 °). Each dot moves independently within its arc, however in general the dots are moving rightwards.

### Experiment 2

#### Design and Apparatus

A series of baseline measures were completed for global motion direction discrimination, contrast sensitivity and orientation discrimination (pre-and post-training assessments). Assessment stimuli were presented on a VIEWPixx/3D 23.6 inch monitor with a display resolution of 1920 x 1080 pixels, with a 120 Hz refresh rate, using a Dell Precision T3610 PC running Windows 7. One pixel subtended 1.6 arc minutes and stimuli were viewed from a distance of 570mm. Head position for testing was stabilised using a chin rest. Following this, all observers undertook five consecutive days of global motion training in one of three spatial frequency groups (broad, high or low). Training stimuli were presented on a 19” monitor with a display resolution of 1980 x 1080 pixels and 60 Hz refresh rate, using a PC running Windows 7. One pixel subtended 1.7 arc minutes. Stimuli were viewed from a distance of 500mm. All stimuli were presented for 1 second.

#### Training Stimuli

##### Global Motion

Broadband stimuli were the same as previously described. For low-frequency stimuli, the elements were circularly symmetric Gabor patches. The standard deviation of the Gaussian window, *σ*, was 30.1 arc minutes, and the spatial frequency of luminance modulation, *f*, was 1 cycle/degree. For each element, the luminance profile was defined as a function of horizontal and vertical position (*x, y*) as:

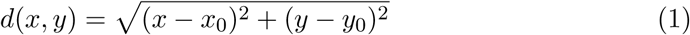

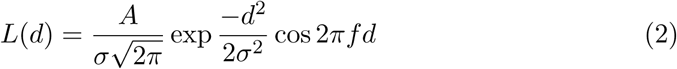

where *x*_0_ and *y*_0_ is the central position of the element, and *A* determines the contrast. Elements for the high frequency stimuli were defined in the same way, but had a standard deviation of 7.48 arc minutes and a spatial frequency, *f*, of 4 cycles/degree. For all stimuli, the spatial frequency and the speed of motion were held constant. Initially all elements were uniformly and randomly distributed within a region of 16.6°x 16.6° on the centre of the screen. A central black fixation dot was presented at all times when stimuli were not being displayed. Examples of the stimuli are shown in see Fig. 2 a-c. Motion was created using the method detailed in Fig. 3. On each frame each dot moved a fixed distance of 8.5 arc minutes.

#### Pre- and Post-training Stimuli

##### Global motion

Stimuli were identical to those described in the training session, with the following exceptions. The standard deviations of the elements were 6.4 arc minutes (broadband), 28.4 arc minutes (low frequency) and 7.0 arc minutes (high frequency). Stimuli were presented within a mid-grey rectangle measuring 15.9°x 15.9°, each element moved a fixed distance of 8 arc minutes.

##### Contrast Sensitivity

Stimuli were Gabor patches, with a spatial frequency of 1 cycle per degree (/°) or 4 cycles/°, presented in the centre of the screen on a mid-grey background, tilted either ±20° (see Fig. 4a&b). The Gaussian envelope of the Gabor stimulus had a standard deviation of 1.1°. 7 levels of contrast (0.05, 0.1, 0.15, 0.175, 0.2, 0.3, 0.4 % Michelson Contrast) were presented.

**Fig 4.**
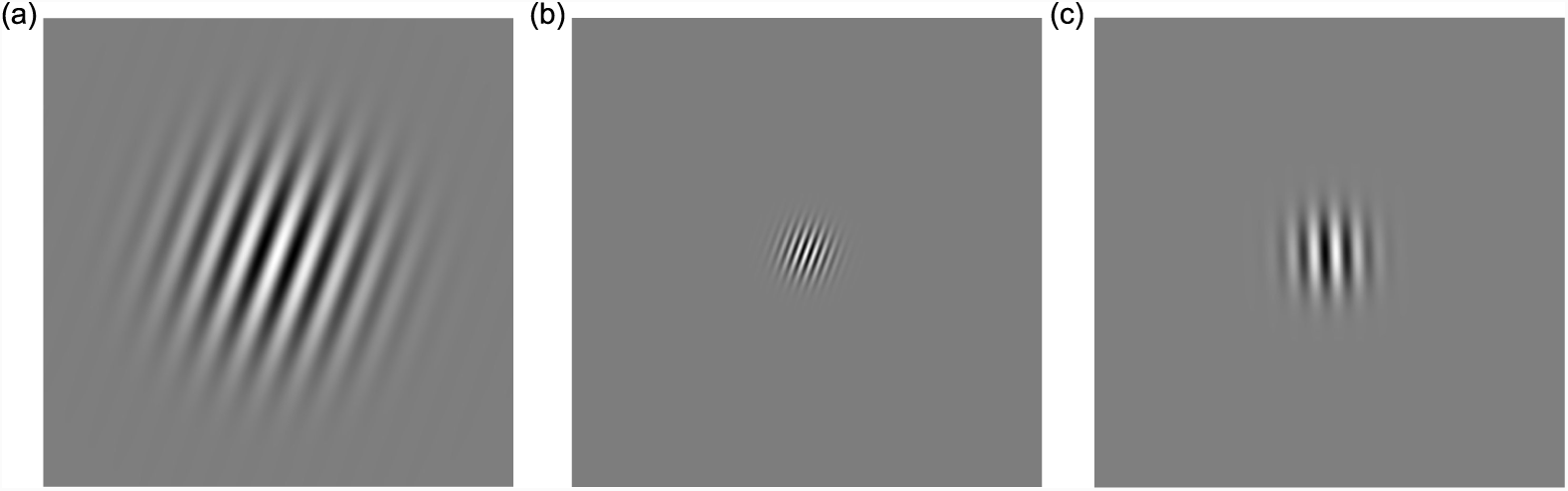
Stimuli: The Gabors used for a) low spatial frequency contrast sensitivity - 1 cycle per degree, b) high spatial frequency-4 cycles per degree, and c) orientation discrimination which is tilted leftwards at 2° with mid spatial frequency-2 cycles per degree

##### Orientation Discrimination

Gabor patches were presented for 100ms on a mid-grey background, measuring 7.9° x 7.9°, and presented 6.6° either above or below the central fixation point. Orientation was centred at zero, at which point lines were vertical. There was a total range of 1.8° difference in tilt, between ± 0.9°, with 7 linearly spaced points within the range. The spatial frequency of the Gabor patch was 1.85 cycles/° and the standard deviation of the Gaussian envelope was 0.2° (see Fig. 4c). Contrast was fixed at 50% Michelson Contrast.

### Procedure

#### Training on Global Motion

For each observer, training was undertaken at one spatial frequency only, totalling 420 trials daily for 5 continuous days. Feedback was provided after each trial.

#### Pre- and Post-Assessments

Measures were taken for global motion (high, broad and low spatial frequency), contrast sensitivity (high and low spatial frequency) and orientation discrimination (angle of tilt from the vertical) of an oriented Gabor patch. Responses were captured on the DataPixx response box for contrast sensitivity, and left and right arrows on the keyboard for global motion and orientation discrimination. The presentation order of trials was randomised for direction and coherence (global motion), orientation and contrast (contrast sensitivity) or orientation (orientation discrimination). There were 20 repetitions for each of the seven levels, for each condition. Testing was performed in a darkened room, before and after training.

#### General Statistical Methods

Moscatelli et al. (2012) [119] proposed using the Generalised Linear Mixed Effects Model (GLMM) for psychophysical data. The GLMM is an extension of the General Linear Model (GLM) that provides a more robust statistical analysis where the data contain irregular response distributions. The GLMM contains both fixed and random effects. The fixed component estimates are the effects of interest, in our study they are a) the day (or session) of testing and b) each level of the stimulus (coherence, contrast or orientation). The modelling of random effects assesses the differences between related groups of data points (such as those from different observers) that allow inference to a larger population [120].

The linear component of the model is given by:

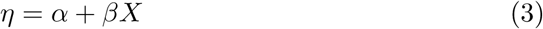

where *α* and *β* are the intercept and the slope of the fixed effects parameters respectively. The link function converts the expected outcome variable (the proportion of correct responses, *p*), to the linear predictor [121]. Here, a logit function was used, adapted to take account of chance performance, such that:

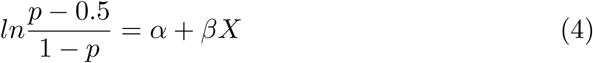

#### Selecting the model

All response data were analysed using a GLMM conducted using Matlab (using the *fitglme* function), the logit link function and REMPL (restricted maximum pseudo likelihood) to estimate the model parameters. Each model included observers as a random effect, and Time and Level and their interaction as fixed effects. Each model was assessed for goodness of fit, using random slopes or intercepts only, or their combination. Models (5a-5d) were compared to test which model produced the lowest model fit statistics [119]. The Akaike information criterion (AIC) is one of the best known criteria to use in selecting a good model, where fitted values are most likely to be true values, thus providing a fit closest to reality [120]. The best regression model will provide the lowest value of AIC.

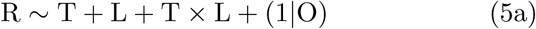

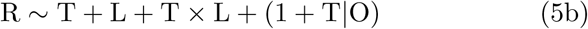

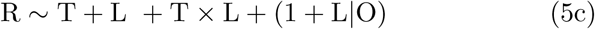

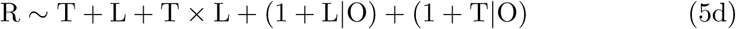

*R* is the observer response (proportion correct) *T* is time, the day or session of testing, *L* is the stimulus level (degree of coherence, level of contrast, or degree of orientation) and *O* is the observer.

#### Interpreting the model

The intercept is the proportion of correct responses when the linear model value is zero. An increase in intercept over time reflects an overall increase in the number of correct responses. The slope relates to how the proportion of correct responses increases with the strength of the stimulus (degree of coherence, level of contrast, or degree of orientation). Overall performance can also be summarised in terms of the 75% threshold, the point at which observers responded correctly on 75% of trials.

## Results

### Feedback and perceptual learning: Experiment 1

To verify the spread of easy and difficult trials across the stimulus levels suggested by Liu et al. [114], accuracy for the first day was evaluated across all observers for each level of the stimulus and is reported in (Table 1). The accuracy across levels confirms that there is an even distribution of easy trials (85% and above) and difficult trials (65% and below) with the balance around 75%.

**Table 1.** Accuracy at each level of the stimulus.

| Stimulus Levels | Coherence ° | Accuracy (%) |
| --- | --- | --- |
| 180 | 180 | 98.1 |
| 300 | 60 | 96.0 |
| 335 | 25 | 80.4 |
| 340 | 20 | 77.7 |
| 345 | 15 | 73.5 |
| 350 | 10 | 64.1 |
| 355 | 5 | 61.5 |

To identify if this distribution of easy and difficult trials was sufficient for learning in the absence of trial-by-trial feedback, we analysed individuals’ daily performance across the two training groups (Feedback or No Feedback). Individual responses from the 10 days’ training were aggregated for each variable; observer, day and level of coherence. The response data were analysed independently for each group (Feedback or No Feedback) using the method outlined above. Response was modelled as a fixed effect of day and coherence and an interaction between these two predictors (see Table 2). For the Feedback Group, there was a significant negative effect of day however there was a significant positive day *×* coherence interaction indicating there was a change in slope across the 10 days. The same analysis was undertaken for the No Feedback group, and found a main effect of day with a significant negative interaction. The no feedback group performed worse after 10 days training (see Fig. 5(b)). Performance over the 10 days is illustrated in Fig. 5(a), where the slope for the feedback group gets steeper over time, day 10 (blue) versus day 1 (red), along with the 75% discrimination threshold, which shifts leftwards, indicating a reduction in threshold between the start and end of training. Conversely, the group trained without feedback performs progressively worse over time and the 75% discrimination threshold shifts rightwards, indicating an increase in threshold.

**Table 2.** Fixed effects parameters for the Pilot Data

| Training | | $\beta$ | SE | tStat | DF | pValue | LCI | UCI |
| --- | --- | --- | --- | --- | --- | --- | --- | --- |
| No Feedback | Intercept (Day 1) | -1.595 | 0.1555 | -10.258 | 836 | 2.456e-23 | -1.90073 | -1.2902 |
|  | Day (Change) | -0.0361 | 0.074 | -0.4886 | 836 | 0.625 | -0.1812 | 0.1089 |
|  | Coherence | 0.07576 | 0.00207 | 36.598 | 836 | 8.65e-176<sup>†</sup> | 0.0717 | 0.0798 |
| | Day × Coherence | -0.00075 | 0.00032 | -2.30953 | 836 | 0.02116<sup>*</sup> | -0.00139 | -0.0001 |
| Feedback | Intercept (Day 1) | -0.6099 | 0.140 | -4.35 | 836 | 1.527e-05 | -0.885 | -0.335 |
|  | Day (Change) | -0.097 | 0.0076 | -12.78 | 836 | 2.78e-34<sup>*</sup> | -0.112 | -0.082 |
|  | Coherence | 0.0292 | 0.0016 | 18.39 | 836 | 1.096e-63<sup>†</sup> | 0.026 | 0.0324 |
| | Day × Coherence | 0.0087 | 0.0004 | 24.3234 | 836 | 3.052e-99<sup>*</sup> | 0.00797 | 0.0094 |

Table notes :<sup>\*</sup> indicates significant p value and <sup>†</sup> performance increases significantly with increasing coherence

**Fig 5.**
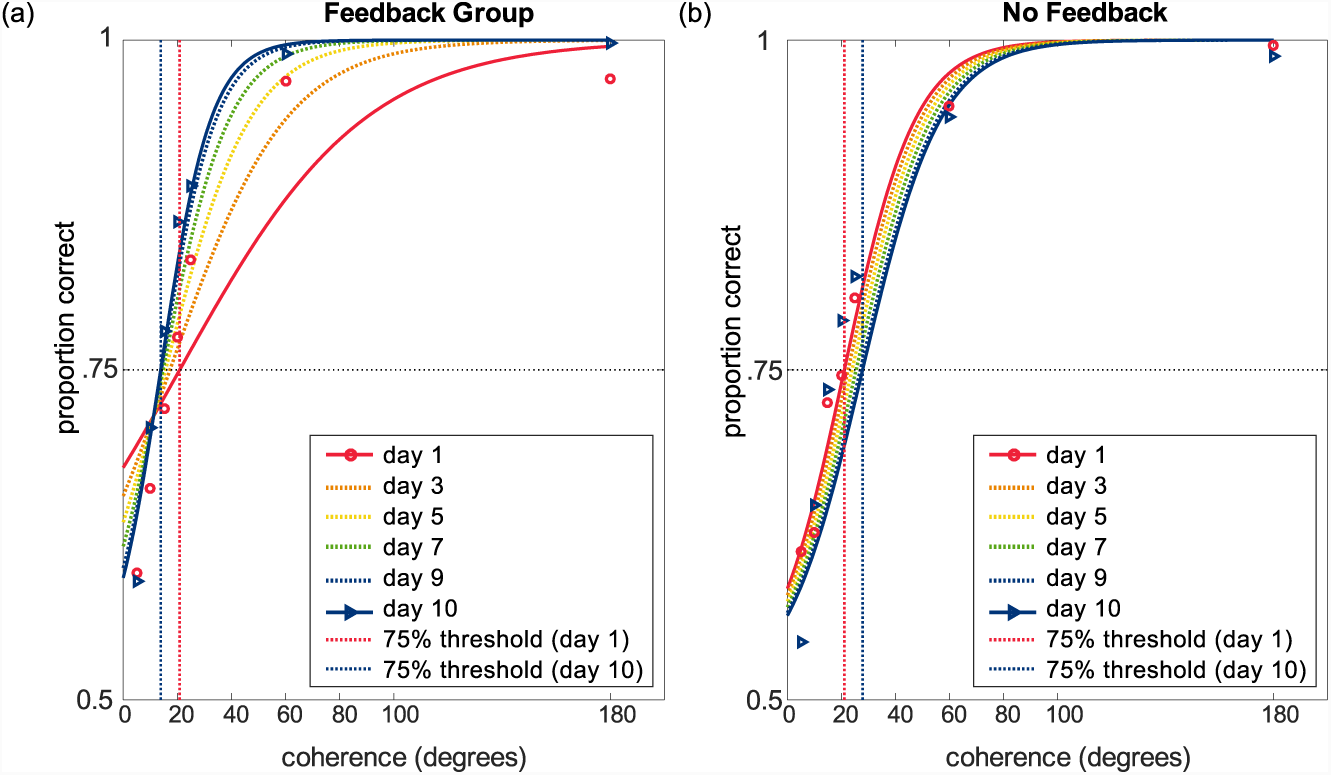
Mean thresholds (in degrees) over 10 days on global motion training for feedback (a) and no feedback (b) groups. The red and blue dotted lines show 75% threshold on days 1 and 10 respectively. 75% threshold decreased for the feedback group, and increased for the no feedback group.

**Fig 6.**
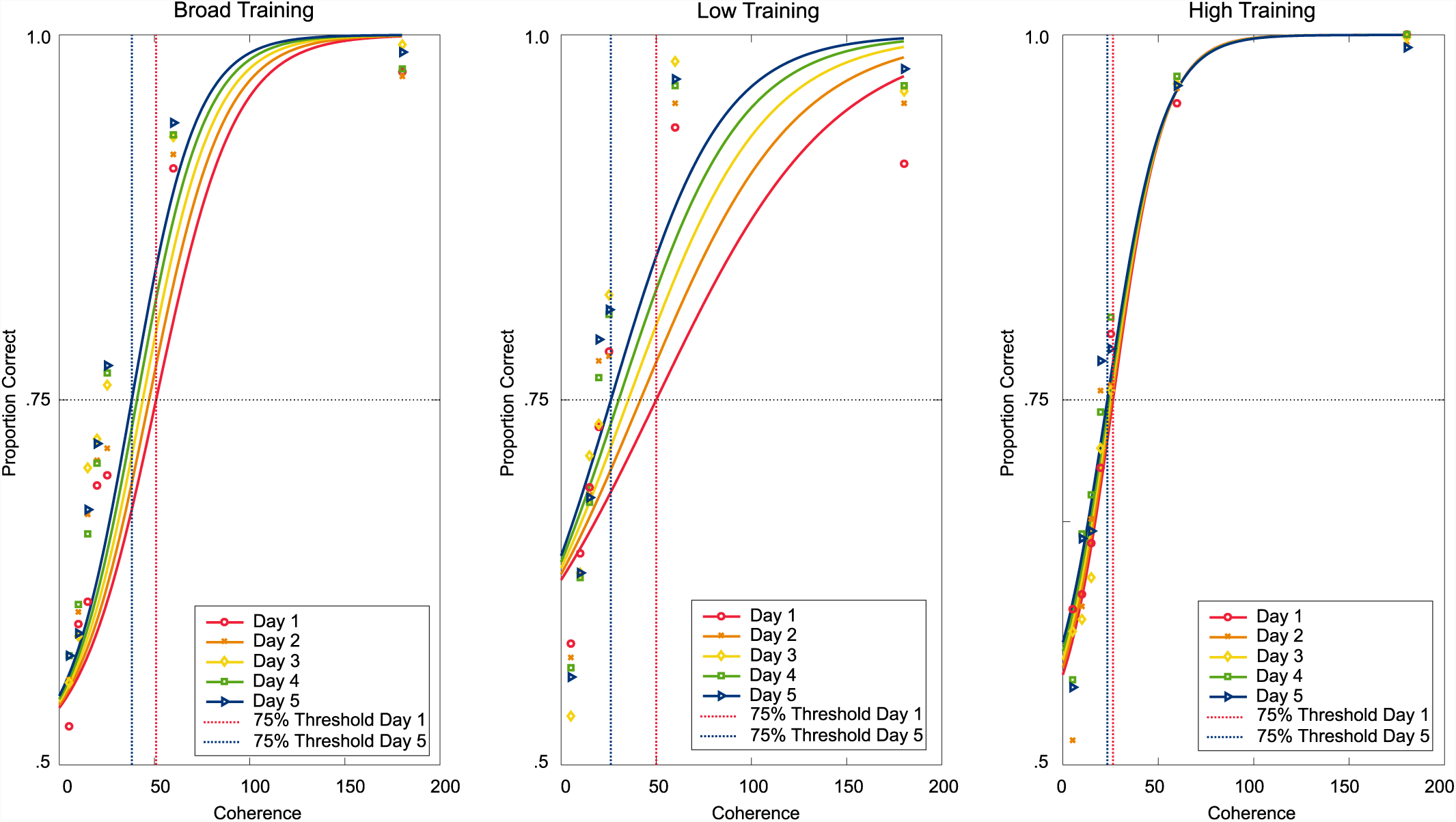
Training results over the 5 days for broad, low and high spatial frequencies. The red and blue dotted lines show 75% threshold on days 1 and 5 respectively.

### Learning and transfer of global motion: Experiment 2

The number of correct responses from the daily training was calculated for each observer, for each day and level of coherence. Training data were analysed independently for the three groups trained on different frequencies (Broad, Low, High) and were modelled as a fixed effect of day and coherence and an interaction between these two predictors.

Results were analysed using the method previously described (see Table 3). Performance improved for the broad and low trained groups and there was a significant positive coherence-by-day interaction found for both conditions. For the group trained with high spatial frequency stimuli there was a significant main effect of day. This shows that there was a significant increase in the total number of correct responses over the five days (see Fig. 3). These results thus show an improvement in performance during the training phase for all three spatial frequency conditions.

**Table 3.**
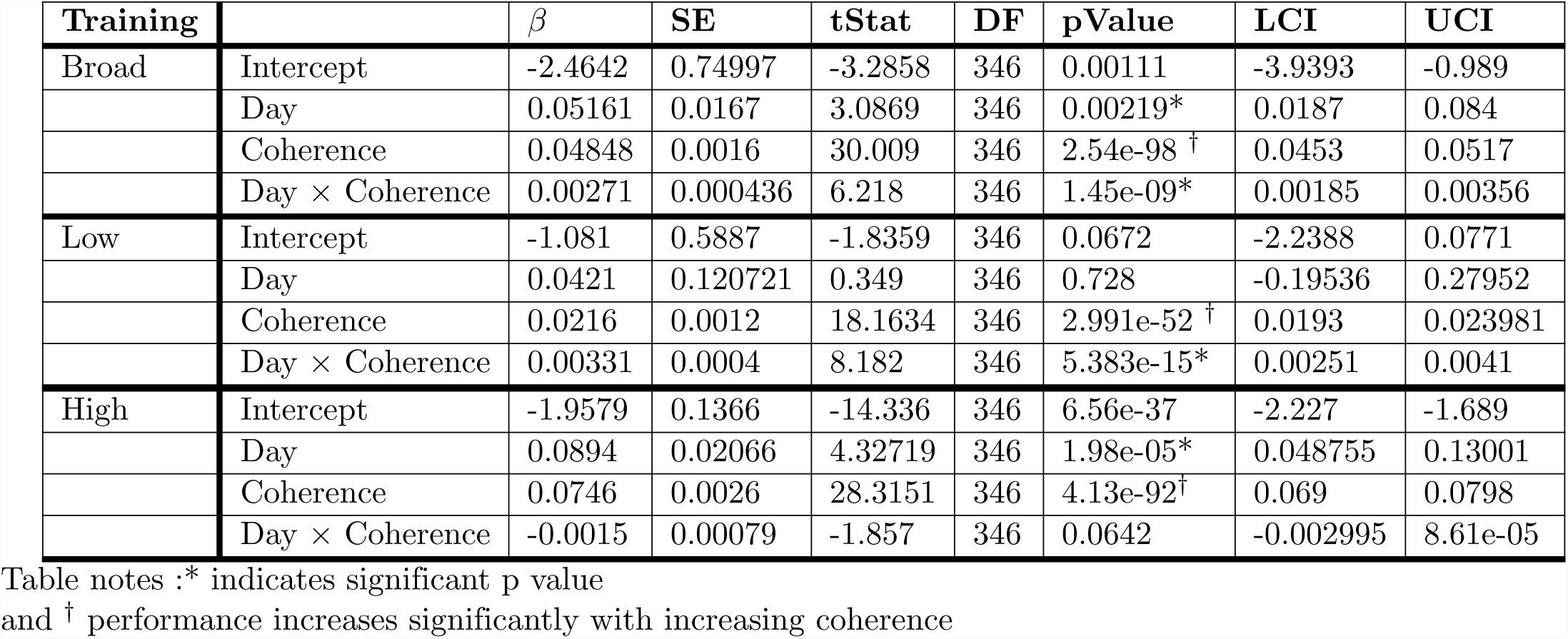
Fixed effects parameters for the Training Data

| Training | | $\beta$ | SE | tStat | DF | pValue | LCI | UCI |
| --- | --- | --- | --- | --- | --- | --- | --- | --- |
| Broad | Intercept | -2.4642 | 0.74997 | -3.2858 | 346 | 0.00111 | -3.9393 | -0.989 |
|  | Day | 0.05161 | 0.0167 | 3.0869 | 346 | 0.00219* | 0.0187 | 0.084 |
|  | Coherence | 0.04848 | 0.0016 | 30.009 | 346 | 2.54e-98<sup>†</sup> | 0.0453 | 0.0517 |
| | Day × Coherence | 0.00271 | 0.000436 | 6.218 | 346 | 1.45e-09* | 0.00185 | 0.00356 |
| Low | Intercept | -1.081 | 0.5887 | -1.8359 | 346 | 0.0672 | -2.2388 | 0.0771 |
|  | Day | 0.0421 | 0.120721 | 0.349 | 346 | 0.728 | -0.19536 | 0.27952 |
|  | Coherence | 0.0216 | 0.0012 | 18.1634 | 346 | 2.991e-52<sup>†</sup> | 0.0193 | 0.023981 |
| | Day × Coherence | 0.00331 | 0.0004 | 8.182 | 346 | 5.383e-15* | 0.00251 | 0.0041 |
| High | Intercept | -1.9579 | 0.1366 | -14.336 | 346 | 6.56e-37 | -2.227 | -1.689 |
|  | Day | 0.0894 | 0.02066 | 4.32719 | 346 | 1.98e-05* | 0.048755 | 0.13001 |
|  | Coherence | 0.0746 | 0.0026 | 28.3151 | 346 | 4.13e-92<sup>†</sup> | 0.069 | 0.0798 |
| | Day × Coherence | -0.0015 | 0.00079 | -1.857 | 346 | 0.0642 | -0.002995 | 8.61e-05 |

Table notes : \* indicates significant p value and <sup>†</sup> performance increases significantly with increasing coherence

### Pre- and Post-Assessment Analysis: Global Motion

Learning is often measured through monitoring performance at a particular threshold, which is expected to shift the psychometric function leftwards if performance is improved, (see Fig. 7(a)). The psychometric function describes performance in terms of accuracy as a function of the strength of the stimulus [122], which are usually positively correlated, and it is expected to reach asymptotic performance at the highest stimulus intensity [123]. Inspection of the pre- and post-assessment data revealed that in some conditions performance did not reach perfect accuracy, asymptoting at a proportion of correct responses that was less than 1; this resulted in a poor psychometric fit of the observer response data using the GLMM. To accommodate this, a nonlinear generalised mixed effects model (NLME) was used to include an additional parameter in order to model variability in the asymptotic performance at high signal levels, as has been applied in other perceptual learning studies [124–126].

**Fig 7.**
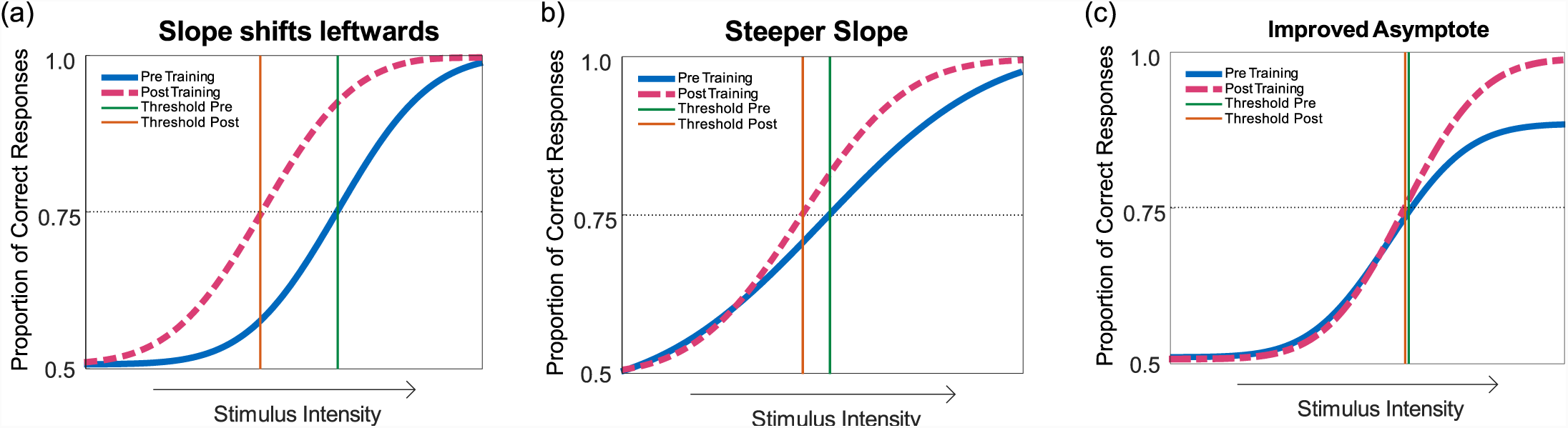
Psychometric functions illustrating the three measures by which the non-linear regression provides evidence of a change. (a) A leftward shift of the function indicates a general increase across stimulus levels, that did not vary as stimulus intensity increased. (b) A steeper slope indicates an increase in the number of correct responses as stimulus intensity increases. (c) An upward shift of the asymptote indicates an increase in performance where stimulus intensity is at its highest. However, in this example there has been no shift of the 75% correct detection threshold.

#### Nonlinear regression analysis

The nonlinear regression provides three measures to assess a change in performance over time. Firstly, like the GLMM a leftward shift in the curve indicates an improvement in threshold (7(a)). An increase in slope indicates an increase in the rate at which performance increases with increase signal level (7(b)). Finally a change in the asymptote indicates a significant change to the performance at the highest level of stimulus intensity (7(c)). These changes are independent aspects of the psychometric function fit, and may not necessarily be congruent. For example, it is possible to obtain an increase in one measure and a decrease (or no change) in another.

Analysis of the pre- and post-assessment data was undertaken using a nonlinear mixed effects regression with the *nlmefit* function in Matlab with the following model:

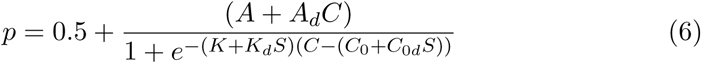

where *p* is the proportion of correct responses, *A* determines the asymptotic level of performance, *K* defines the slope and *C*_0_ defines the threshold. *d* is the post-training change in performance, where *A*_*d*_, *K*_*d*_ and *C*_0 *d*_ determine the change in asymptote, slope and threshold respectively. *C* is the coherence level, and *S* is a dummy variable, taking on values of 0 (pre) or 1 (post) training sessions.

95% confidence limits were calculated using parametric bootstrap for 1000 simulated experiments, with the same number of simulated observers and repetitions as the actual experiment. Results for each condition are plotted in Figs. 8 to 10.

**Fig 8.**
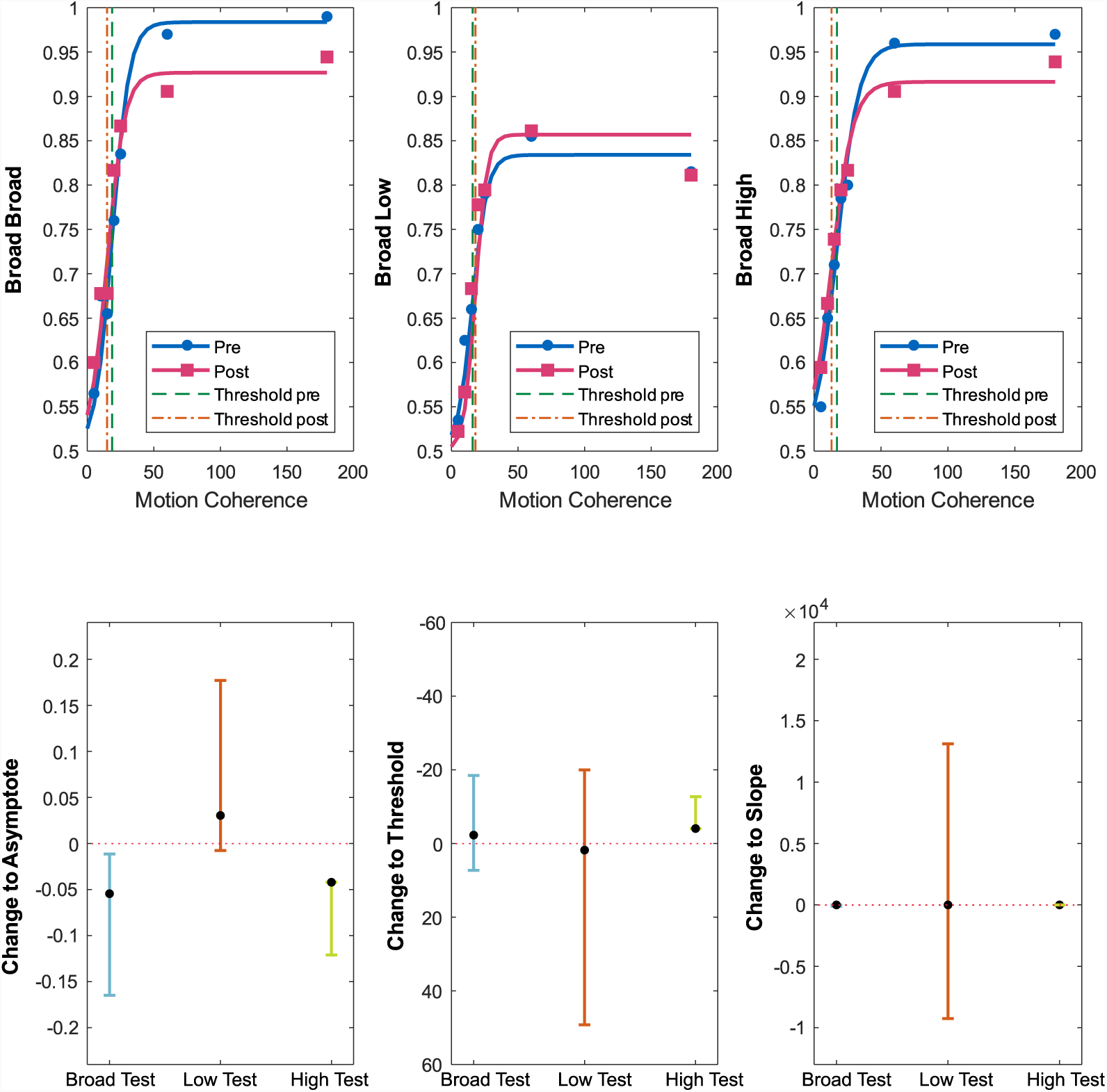
**Top:** Assessment results for the Broad Frequency trained condition on Broad, Low and High spatial frequency tests. Performance is presented as the proportion of correct responses, where blue circles show the results for the pre-assessment, and pink squares for the post-assessment. 75% thresholds for pre and post are indicated by the green and orange vertical dotted lines respectively. **Bottom:** Change statistics for the asymptote, threshold and slope, respectively. Plots show the median performance and the 95% confidence intervals for the change in performance between pre- and post-assessments. The red horizontal line at zero represents no change, confidence intervals crossing the zero line reflect no significant improvement. Points above the reference line show an improvement in performance and those below reflect a decrease in performance.

**Fig 9.**
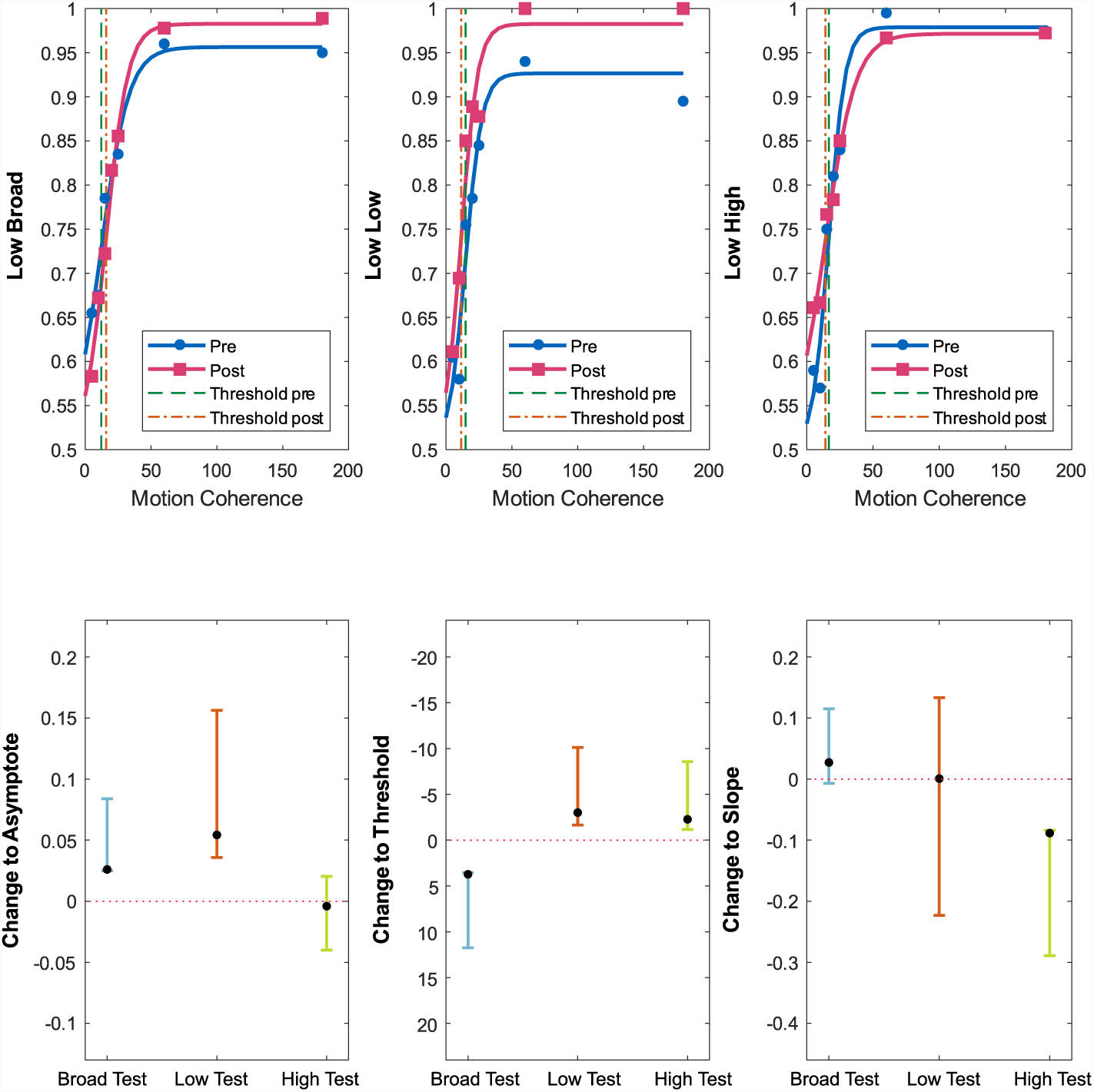
**Top:** Assessment results for the Low Frequency trained condition on Broad, Low and High spatial frequency tests. Performance is presented as proportion of correct responses, where blue (circles) show the results for the pre-assessment, and pink (squares) the post-assessment. 75% thresholds for pre- and post-assessments are indicated by the green and orange vertical dotted lines respectively. **Bottom:** Change statistics for the asymptote, threshold and slope respectively. Plots show the median performance and the 95% confidence intervals for the change in performance between pre- and post-assessments. The red horizontal line at zero represents no change, confidence intervals crossing the zero line reflect no significant improvement. Points above the reference line show an improvement in performance and those below reflect a decrease in performance.

**Fig 10.**
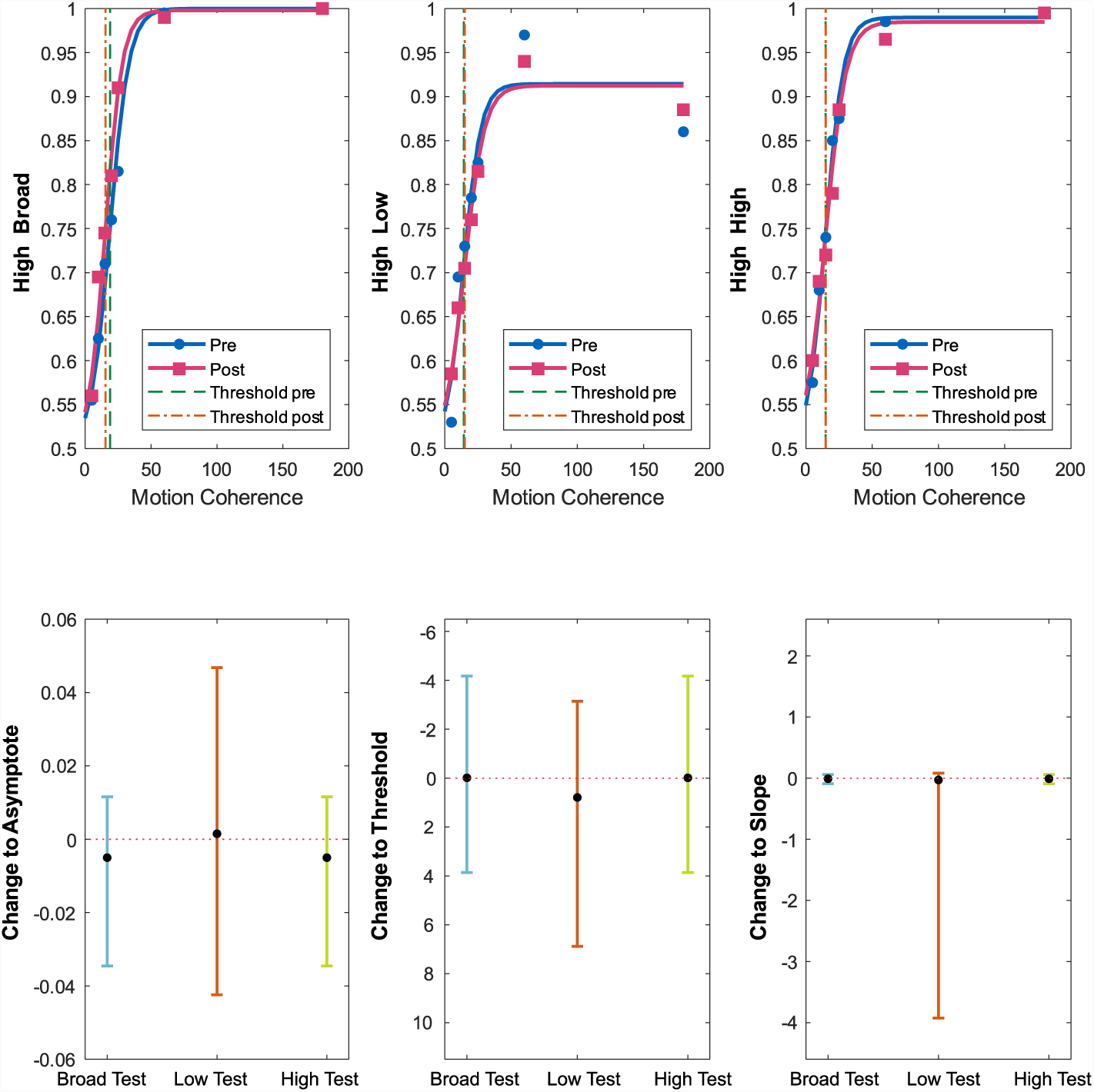
**Top:** Assessment results for the High Frequency trained condition on Broad, Low and High spatial frequency tests. Performance is presented as proportion of correct responses, where blue (circles) show the results for the pre-assessment, and pink (squares) the post-assessment. 75% thresholds for pre and post are indicated by the green and orange vertical dotted lines respectively. **Bottom:** Change statistics for the asymptote, threshold and slope respectively. Plots show the median performance and the 95% confidence intervals for the change in performance between pre- and post-assessments. The red horizontal line at zero represents no change, confidence intervals crossing the zero line reflect no significant improvement. Points above the reference line show an improvement in performance and those below reflect a decrease in performance.

### Predictions

We predicted that should transfer occur cortically, this would be located at the global motion processing level, and transfer across frequencies would most likely occur for stimuli with frequency properties that reflect the broadband low-pass frequency tuning of global motion detectors (see Fig 11(a)). Firstly, we predicted no condition would show transfer to contrast sensitivity, as these are processed in different cortical locations. For global motion, we predicted;

**Fig 11.**
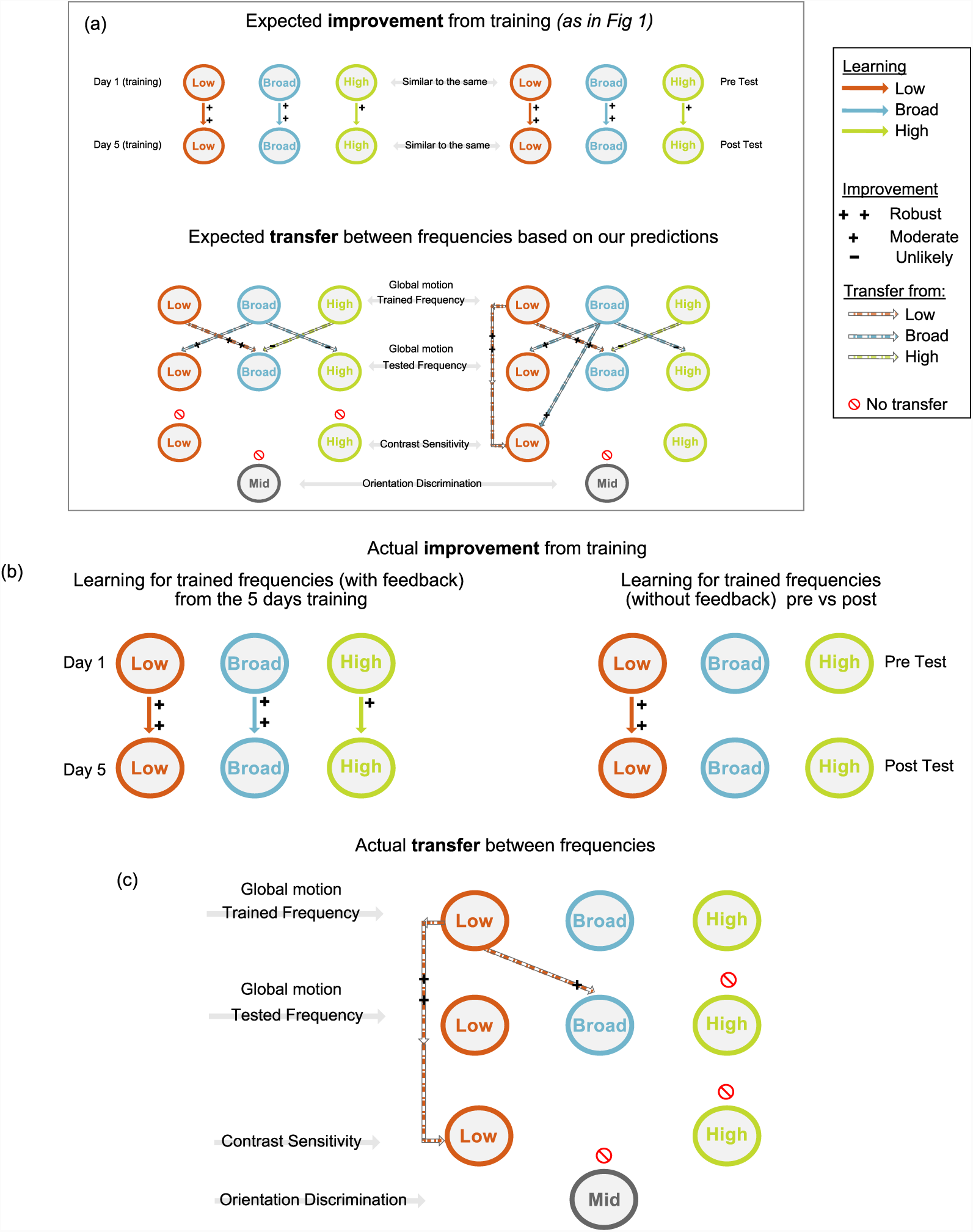
(a) Predictions of learning and transfer as shown in Fig. 1. We predicted a) that if transfer was local to the processing mechanism transfer would occur for global motion conditions with shared spatial frequency properties. (b) Performance improved for all groups during the five days training, with robust improvement for the low and broad trained groups, and moderate improvement for the high frequency group. However, at the post-assessment stage, when tested without the presence of trial by trial feedback, neither the broad nor high groups reflected a change from the pre-assessment measures. Improvement was restricted to the low trained group.(c) When assessing transfer to untrained frequencies, only the low spatial frequency group showed evidence of transfer, and performance improved for broad frequency motion. In addition, the low spatial frequency trained group was the only group to show an improvement on contrast sensitivity and this was exclusive to the low frequency contrast condition.

- For the broad frequency trained condition; (i) robust learning on their own trained frequency (ii) moderate transfer to low spatial frequency global motion, and (iii) an unlikely but possible transfer to the high frequency condition.
- For the low frequency trained condition; (i) robust learning on their own trained frequency and (ii) robust transfer to broad spatial frequency global motion.
- For the high frequency trained condition we predicted; (i) moderate learning on their own trained frequency and (ii) an unlikely but possible transfer to the broad frequency condition.

Conversely, should transfer occur as a result of backwards generalisation using the re-entrant connections from global motion processing areas to V1 we would predict that transfer would reflect the pooling of spatial frequencies and likely attenuation of high spatial frequencies.

- For the broad frequency trained condition; (i) robust learning on their own trained frequency (ii) moderate transfer to low spatial frequency global motion, (iii) a possible but unlikely transfer to high spatial frequency global motion and (iv) a moderate transfer to low spatial frequency contrast sensitivity
- For the low frequency trained condition; (i) robust learning on their own trained frequency and (ii) robust transfer to broad spatial frequency global motion, and (iii) robust transfer to low spatial frequency contrast sensitivity
- For the high frequency trained condition we predicted; (i) moderate learning on their own trained frequency and (ii) an unlikely but possible transfer to the broad frequency condition, and (iii) no transfer to any frequency of contrast sensitivity

### Transfer to global motion: broad-frequency trained group

We had predicted robust to moderate improvement for the broad trained group, however this group showed no consistent transfer to trained or untrained conditions (see Fig. 8). The post-assessment on the **trained task** found no significant change in slope *(proportion correct as a function of frequency)* or threshold *(coherence required to obtain 75 % correct)*, and a significantly lower asymptote *(proportion of correct responses at the highest coherence).* Assessing transfer to **untrained tasks**, there was no significant change for any measure (intercept, slope or asymptote) for the **low frequency** test. Finally for the **high frequency** test, there was no change in slope, a significant reduction in asymptote (indicating worse performance at higher coherence), and a small but significant reduction in threshold. The increase in the proportion of correct responses at the lower coherence levels was the only significant improvement for this group, however neither of the other two measures were consistent with an improvement.

### Transfer to global motion: low-frequency trained group

The low-frequency trained group was the only condition to exhibit transfer to trained and untrained motion conditions (see Fig. 9). In the post-assessment on the **trained task**, while there was no significant change to the slope *(proportion correct as a function of frequency)* there was significant reduction in threshold *(coherence required to obtain 75% correct)*, and a significantly higher asymptote *(proportion of correct responses at the highest coherence).* Assessing transfer to **untrained tasks**, there was a modest improvement for the **broad frequency** test reflected by the was a significant increase in asymptotic performance. However, there was an increase in threshold and no change to the slope. This shows reduced sensitivity at low signal levels, but an increase is performance at higher levels. Finally, the **high frequency** test stimuli show a small reduction in threshold but a significantly shallower slope. Suggestive of increased performance at lower coherence but decreased performance at higher stimulus coherence. There was no change in asymptotic performance, and overall no evidence of transfer.

### Transfer to global motion: high-frequency trained group

The group trained with high spatial frequency stimuli showed no transfer to any trained or untrained motion condition for any of the three measures (see Fig. 10).

### Transfer to global motion: Participant level variability

To investigate the lack of transfer to the trained task, a participant level plot (see Fig.12) illustrates the change in number of correct responses between pre- and post-training. This was calculated by subtracting the pre-assessment scores from the post-assessment scores. Changes above the dotted line at 0 show improved performance, while those below the line show worsened performance. The spread of scores for the broad/broad and broad/high condition show that one observer performed particularly poorly at higher coherence levels, but not persistently so. Therefore, the lack of overall improvement for the broad trained group is unlikely to be as a result of an outlier. However, the significantly worse performance in asymptotic performance may, to some degree, be as a result of this one individual.

### Transfer to global motion: Summary

In summary, only the low frequency trained group provided reliable evidence for transfer to trained and untrained motion conditions, with a moderate transfer to the broadband frequency condition.

**Fig 12.**
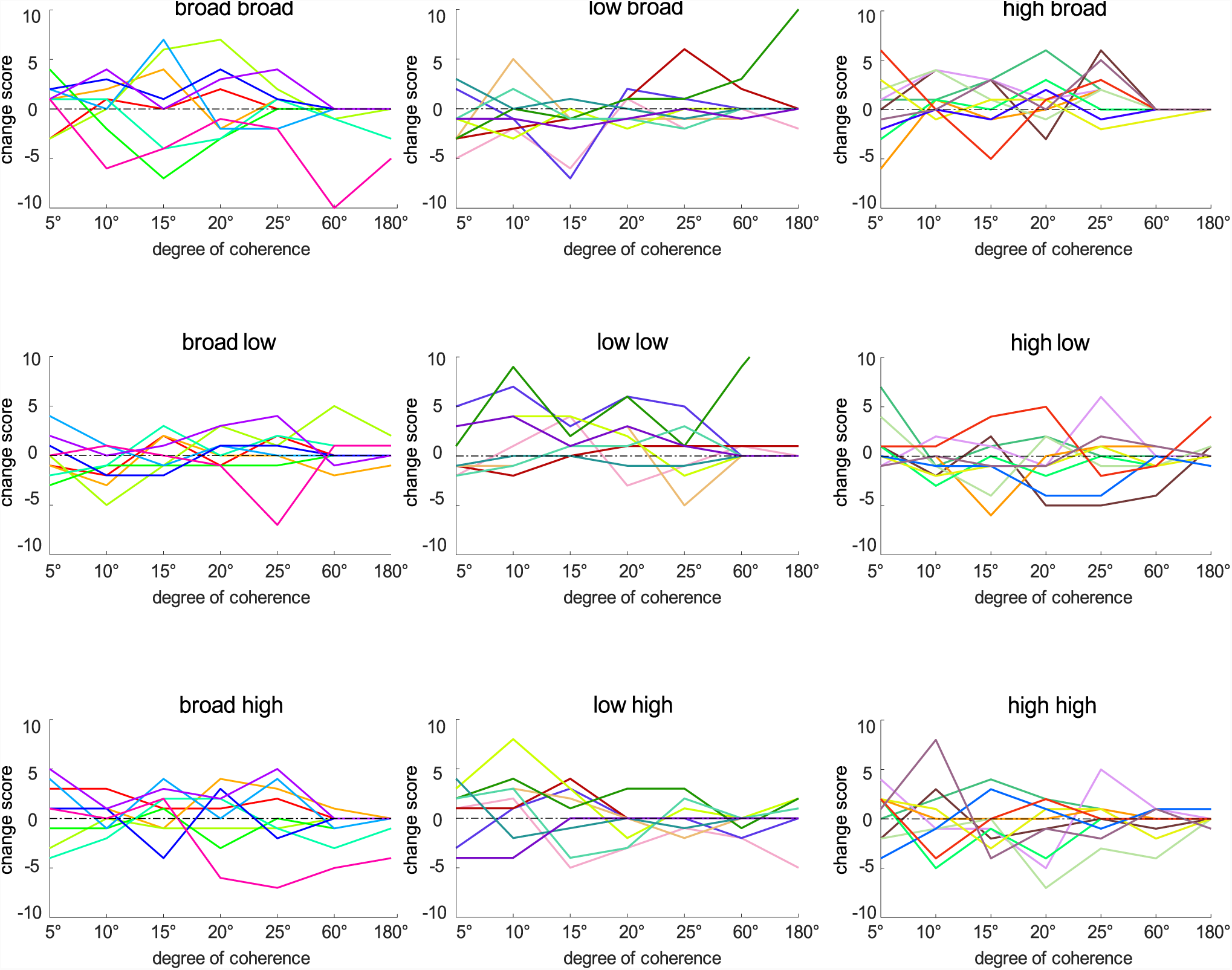
Change scores (the difference between the number correct on the pre- and post-assessments) are plotted for each individual. Each observer is illustrated by a colour grouped by training and test condition (broad, low and high spatial frequency motion). Of particular interest is the broad trained group, which had a decline in performance post training. One observer performed worse at higher coherence levels after training, which explains the decline in asymptotic performance for the broad trained group. However, the overall spread of data suggest that on average post-assessment results did not differ significantly.

### Transfer to contrast sensitivity

We predicted that, should contrast sensitivity improve as a result of training on global motion it could only occur as a result of backward generalisation, as the two tasks are processed in cortically separate locations. We predicted transfer would be to limited conditions containing low frequencies, namely the broad and low trained groups.

Data were analysed for all but the same 2 observers who did not complete the final session using the generalised linear effects model previously outlined. The measures of analysis are the intercept and slope, and an interaction between the two predictors.

**Table 4.**
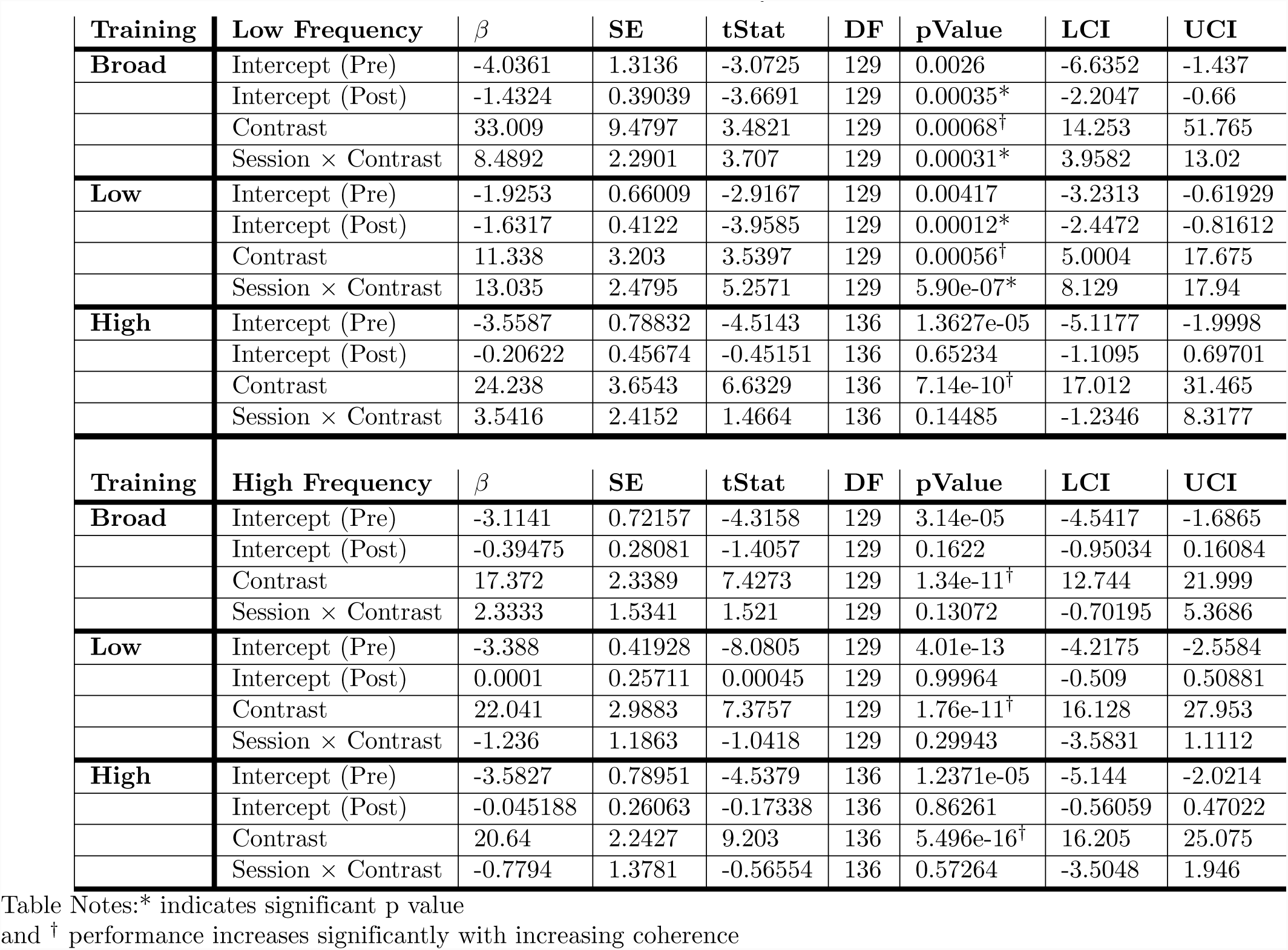
Contrast Sensitivity Table

| Training | Low Frequency | $\beta$ | SE | tStat | DF | pValue | LCI | UCI |
| --- | --- | --- | --- | --- | --- | --- | --- | --- |
| **Broad** | Intercept (Pre) | -4.0361 | 1.3136 | -3.0725 | 129 | 0.0026 | -6.6352 | -1.437 |
|  | Intercept (Post) | -1.4324 | 0.39039 | -3.6691 | 129 | 0.00035* | -2.2047 | -0.66 |
|  | Contrast | 33.009 | 9.4797 | 3.4821 | 129 | 0.00068<sup>†</sup> | 14.253 | 51.765 |
| | Session × Contrast | 8.4892 | 2.2901 | 3.707 | 129 | 0.00031* | 3.9582 | 13.02 |
| **Low** | Intercept (Pre) | -1.9253 | 0.66009 | -2.9167 | 129 | 0.00417 | -3.2313 | -0.61929 |
|  | Intercept (Post) | -1.6317 | 0.4122 | -3.9585 | 129 | 0.00012* | -2.4472 | -0.81612 |
|  | Contrast | 11.338 | 3.203 | 3.5397 | 129 | 0.00056<sup>†</sup> | 5.0004 | 17.675 |
| | Session × Contrast | 13.035 | 2.4795 | 5.2571 | 129 | 5.90e-07* | 8.129 | 17.94 |
| **High** | Intercept (Pre) | -3.5587 | 0.78832 | -4.5143 | 136 | 1.3627e-05 | -5.1177 | -1.9998 |
|  | Intercept (Post) | -0.20622 | 0.45674 | -0.45151 | 136 | 0.65234 | -1.1095 | 0.69701 |
|  | Contrast | 24.238 | 3.6543 | 6.6329 | 136 | 7.14e-10<sup>†</sup> | 17.012 | 31.465 |
| | Session × Contrast | 3.5416 | 2.4152 | 1.4664 | 136 | 0.14485 | -1.2346 | 8.3177 |
| Training | High Frequency | $\beta$ | SE | tStat | DF | pValue | LCI | UCI |
| **Broad** | Intercept (Pre) | -3.1141 | 0.72157 | -4.3158 | 129 | 3.14e-05 | -4.5417 | -1.6865 |
|  | Intercept (Post) | -0.39475 | 0.28081 | -1.4057 | 129 | 0.1622 | -0.95034 | 0.16084 |
|  | Contrast | 17.372 | 2.3389 | 7.4273 | 129 | 1.34e-11<sup>†</sup> | 12.744 | 21.999 |
| | Session × Contrast | 2.3333 | 1.5341 | 1.521 | 129 | 0.13072 | -0.70195 | 5.3686 |
| **Low** | Intercept (Pre) | -3.388 | 0.41928 | -8.0805 | 129 | 4.01e-13 | -4.2175 | -2.5584 |
|  | Intercept (Post) | 0.0001 | 0.25711 | 0.00045 | 129 | 0.99964 | -0.509 | 0.50881 |
|  | Contrast | 22.041 | 2.9883 | 7.3757 | 129 | 1.76e-11<sup>†</sup> | 16.128 | 27.953 |
| | Session × Contrast | -1.236 | 1.1863 | -1.0418 | 129 | 0.29943 | -3.5831 | 1.1112 |
| **High** | Intercept (Pre) | -3.5827 | 0.78951 | -4.5379 | 136 | 1.2371e-05 | -5.144 | -2.0214 |
|  | Intercept (Post) | -0.045188 | 0.26063 | -0.17338 | 136 | 0.86261 | -0.56059 | 0.47022 |
|  | Contrast | 20.64 | 2.2427 | 9.203 | 136 | 5.496e-16<sup>†</sup> | 16.205 | 25.075 |
| | Session × Contrast | -0.7794 | 1.3781 | -0.56554 | 136 | 0.57264 | -3.5048 | 1.946 |

Table Notes:\* indicates significant p value and <sup>†</sup> performance increases significantly with increasing coherence

### Broad-frequency trained group

For low spatial frequency contrast there was a a significant decrease in intercept *(fewer correct responses at the lowest coherence)*, but significant increase in slope *(an increase in the proportion of correct responses as a function of frequency)*. Because of this ambiguity we plotted 75% thresholds for pre and post performance (see Fig. 13 top left). Post-assessment thresholds shifted rightwards suggesting worse performance overall.

**Fig 13.**
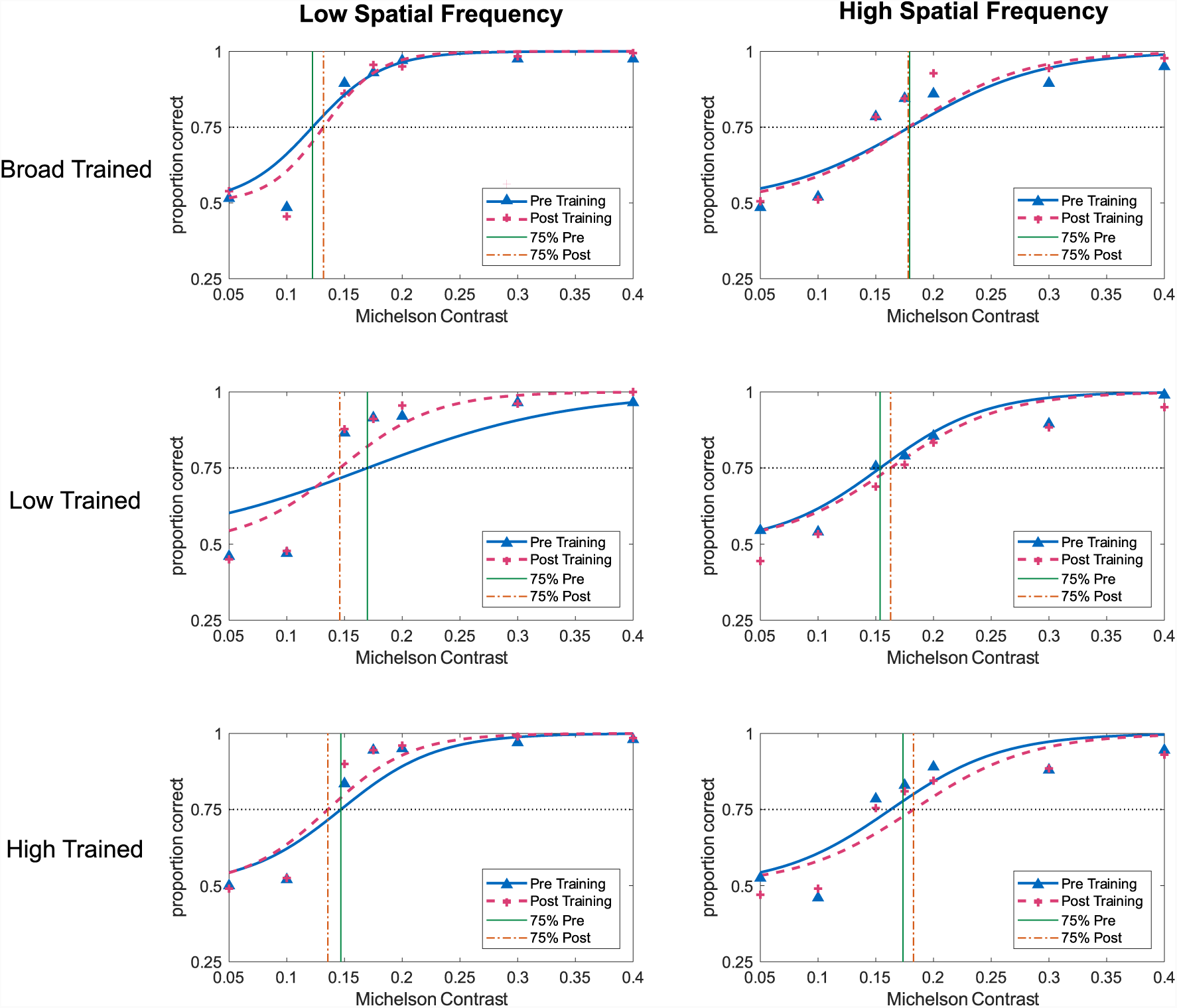
Contrast sensitivity results for Broad (top row), Low (middle) and High (bottom) trained groups for pre- and post-assessments. Low spatial frequencies are shown in the left column and high spatial frequencies on the right. Performance is presented as the proportion of correct responses as a function of Michelson Contrast. Blue (triangles) show the results for the pre-assessment, and pink (crosses) the post-assessment. 75% thresholds for pre and post are indicated by the green and orange vertical lines respectively.

### Low-frequency trained group

The group trained with low frequency global motion showed a significant decrease in intercept, and a significant increase in slope. Again, we plotted 75% thresholds for pre and post performance (Fig. 13 left middle). In this instance the steeper slope, and leftward shift of the threshold, suggests improvement overall.

### High-frequency trained group

The high frequency trained group showed no significant change for any condition (see Fig. 13 bottom left).

### Pre and Post Analysis: Orientation Discrimination

This task was added as an untrained control, for which we predicted no improvement, as no orientation information was present in any of the training stimuli. Data were analysed for all but 3 observers who did not complete the final session, using the generalised linear effects model previously outlined. The measures of analysis are the intercept and slope, and an interaction between the two variables. For analysis, the orientation levels were converted to absolute values. Results are plotted in Fig. 14, and the fixed effects are listed in tab 5. Neither the broad- or low-frequency trained group showed any significant change on this task. However, the high spatial frequency trained group showed a significant decrease in intercept over session, and a significant increase in slope following training. However, the 75% threshold at post-assessment shift rightwards, suggesting worse performance overall.

**Fig 14.**
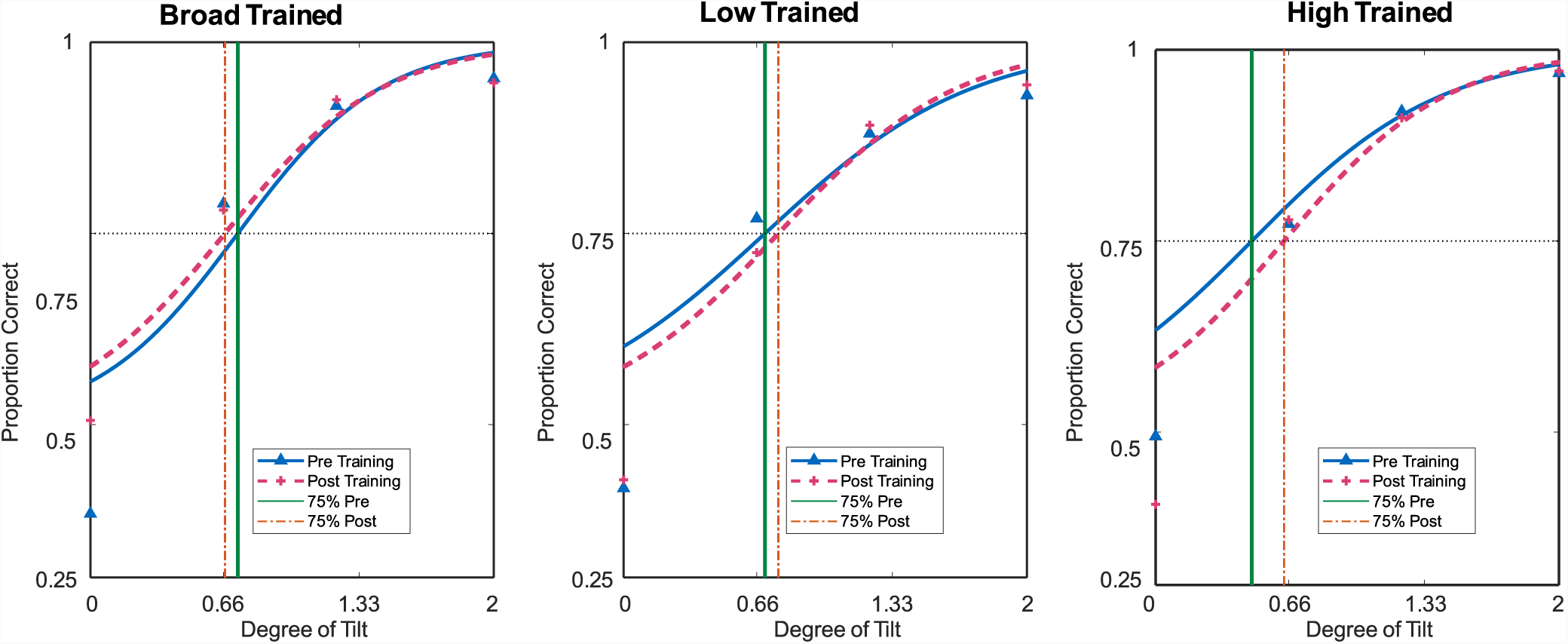
Orientation discrimination results for all trained groups (broad, low and high) for pre- and post-assessments. Performance is presented as proportion of correct responses, where blue (triangles) show the results for the pre-assessment, and pink (crosses) the post-assessment. 75% thresholds for pre and post are indicated by the green and orange vertical lines respectively.

## Discussion

Damage to the visual cortex, as a result of stroke or other brain injury can result in dramatic changes to connectivity between areas. It has been proposed that perceptual training may play a role in strengthening existing neural pathways and may even create novel connections [1–7]. While specificity is a limitation to the effectiveness of perceptual learning as a tool for therapy, there is evidence that this specificity is reduced for visually impaired populations [127–132]. Huxlin et al. (2009) [5] proposed that using global motion may tap into “islands of activity” within V1 through the feedback connections from V5 to V1. However, it is still unclear how much the brain is able to compensate for damage, and whether recovery involves building new connections or changes in the functional connectivity within the existing pathways [133]. Understanding how the cortical hierarchy is organised in terms of the nature of the feedforward and re-entrant pathways is central to developing theories of perception [134].

**Table 5.** Fixed Effects table for Orientation Discrimination

| Training | | $\beta$ | SE | tStat | DF | pValue | LCI | UCI |
| --- | --- | --- | --- | --- | --- | --- | --- | --- |
| Broad | Intercept (Pre) | -1.9841 | 0.79477 | -2.4964 | 284 | 0.013 | -3.5485 | -0.41967 |
|  | Intercept (Post) | 0.02189 | 0.55128 | 0.039723 | 284 | 0.96834 | -1.0632 | 1.107 |
|  | Orientation | 2.8444 | 0.35047 | 8.1161 | 284 | 1.47e-14<sup>†</sup> | 2.1546 | 3.5343 |
| | Session × Orientation | 0.0025 | 0.10152 | 0.0245 | 284 | 0.981 | -0.197 | 0.2023 |
| Low | Intercept (Pre) | -1.5733 | 0.78192 | -2.0121 | 284 | 0.045 | -3.1124 | -0.034191 |
|  | Intercept (Post) | -0.38185 | 0.58052 | -0.65778 | 284 | 0.511 | -1.5245 | 0.76081 |
|  | Orientation | 2.3483 | 0.29959 | 7.8381 | 284 | 9.25e-14<sup>†</sup> | 1.7586 | 2.938 |
| | Session × Orientation | 0.18935 | 0.097122 | 1.9496 | 284 | 0.0522 | -0.0018 | 0.38052 |
| High | Intercept (Pre) | -1.1825 | 0.37792 | -3.1289 | 284 | 0.00194 | -1.9263 | -0.43858 |
|  | Intercept (Post) | -0.47101 | 0.1002 | -4.7005 | 284 | 4.05e-06 | -0.66824 | -0.27377 |
|  | Orientation | 2.3751 | 0.33682 | 7.0516 | 284 | 1.35e-11<sup>†</sup> | 1.7121 | 3.0381 |
| | Session × Orientation | 0.28973 | 0.08728 | 3.3196 | 284 | 0.001* | 0.11794 | 0.46153 |

Table notes : <sup>†</sup> performance increases significantly with increasing coherence

### Feedback and Perceptual Learning: Experiment 1

The purpose of the first experiment was to establish the necessity of performance feedback when undertaking a period of training on a task to discriminate the direction of global motion. Fahle and Edelman (1993) [28] predicted that internal reinforcement could act as the teaching signal when performance feedback was absent and when the confidence is high [34, 135]. Therefore, perceptual learning should occur when training procedures include a mixture of easy and difficult trials. This was the case for the study conducted by Liu et al. (2012) [114] where learning occurred for easy and difficult trials without feedback. For this specific task, consistent with Seitz et al. (2006) [136], our results found that feedback was a requirement for learning. After ten days of training the group who received feedback improved significantly, while the performance for the no feedback group deteriorated.

Our predictions were based on the findings of Liu et al. (2012) [114] who demonstrated that interleaving high accuracy (easy) trials and low accuracy (difficult) trials, resulted in perceptual learning without the need for feedback. Liu et al. (2012) [114] explain their findings using the Augmented Hebbian Re-weighting Model (AHRM) [21]. When external feedback is provided, the post-synaptic activation is shifted further in the correct direction, enforcing appropriate weight changes in the decision unit. However, when external feedback is absent the model uses the observer’s internal response. In this situation learning is dependent on the level of difficulty of the task, and uses the observer’s internal confidence to update the weights [114]. Where a task is easy, the weights still move, on average, in the correct direction [137]. For a difficult task, the neural signal is weak and a clear indication of the appropriate changes that are required is absent. As a result the process of updating the decision weights is ineffective, and learning does not occur.

The results of our study showed that learning did not occur without feedback, even when easy trials were presented. While our task spanned the 65-85% accuracy range used by Liu et al. (2012) [114], there were some differences between the studies. Firstly, Liu et al. (2012) used an adaptive staircase to track performance and stimuli were presented at either 85% or 65% accuracy, and at no other levels, throughout the experiment. Ours, on the other hand, did not include an adaptive staircase. Stimuli were randomly presented using MOCS. This method involves random stimulus selection from a predefined range of stimulus magnitudes [138]. In our experiment observers were randomly presented 1 of 7 threshold levels. Interleaving these levels may influence how decision weights are updated and reduce the observer’s confidence in their judgements, for example by disrupting observers’ meta-cognitive judgements of perceptual confidence [139] and thus their ability to selectively weight high-confidence trials in perceptual learning [34].

Our findings bear some similarity to those found by Seitz et al. (2006) [136], who also used MOCS. They found that learning did not occur without feedback, even when easy trials were presented. Observers trained on one of two tasks, either to discriminate the direction of low luminance motion stimuli, or to discriminate the orientation of a bar that had been masked in spatial noise. After training, both groups who had received feedback showed an improvement, while the groups without feedback did not improve. Seitz et al. (2006) [136] note that many of the experiments investigating perceptual learning use adaptive staircase procedures, and even though easy trials were present, this difference may contribute to the difference in findings. Seitz et al. (2006) [136] propose that interleaving easy and difficult trials within the staircase may allow for “better bootstrapping” from easy to hard, compared to trials that are randomly presented.

A second difference between the study conducted by Liu et al. (2012) and ours, was the type of task used. We used a global motion coherence task and not an orientation discrimination task [114]. These tasks are known to be processed at different levels of the visual hierarchy. Learning without feedback has been obtained for local motion tasks. Ball and Sekuler (1982) [15] found that observers did not need feedback to improve on their direction discrimination task. In their study, observers were required to make a same/different judgement for two rapidly presented trials. In the “same” trials, motion took the same direction, and in the “different” trials the direction of motion varied by 3°. However, while the no feedback group did not receive trial-by-trial response feedback, they were rewarded with two cents for a correct response and had one cent deducted for each incorrect response, which may be construed as end of block feedback [136], which has been found to be as effective as trial-by-trial feedback [113].

Learning without feedback has also been found for global motion coherence tasks [78, 140]. However, none of these examples used the equivalent noise coherence task, but rather used the ratio of signal-to-noise coherence task. Levi et al. [75], using an equivalent noise method of global motion, also found perceptual learning, however they used an adaptive staircase for training paired with trial-by-trial feedback. When investigating task-irrelevant learning, Watanabe et al. (2002) [141] found that task-irrelevant local motion improved passively, but did not find the same for task-irrelevant global motion. They suggest that this is indicative of the lower levels of the visual hierarchy being more receptive to modification, when attention is limited. It was unexpected for the group trained without feedback that performance would deteriorate, and highlights interesting questions about motivation and performance over time as confidence and attention declines. The results of this study ultimately provide us with evidence that to obtain learning during training, using the equivalent noise motion coherence used by Levi et al. (2015) [75], and presenting stimuli randomly using MOCS, our procedure should include trial-by-trial feedback during the training process. Furthermore, since learning did not occur without feedback, no feedback would be provided during the pre- and post-assessment phases.

### Learning and Transfer of Global Motion: Experiment 2

Having established the necessity of feedback for learning, the objective of the main study was to investigate the specificity of spatial frequency tuning in perceptual learning for global motion. We trained three groups of observers on a global motion task with stimuli tuned to three different spatial frequency ranges (Broad, Low and High) and performed pre- and post-training assessments for all frequencies and high and low spatial frequency contrast detection.

### Summary of Training

Our training results demonstrated that after the five day training period on a frequency specific global motion coherence task with trial-by-trial feedback, there was significant improvement in performance for all trained groups.

### Pre- and Post-Assessment predictions

Based on the evidence for the frequency tuning of V5 [79, 80, 96, 104], we predicted that we might have obtained greater improvements (and transfer) between broad and low spatial frequencies, but limited or no transfer for the high spatial frequency trained group. Furthermore, we predicted that should transfer occur as a result of the reweighting of the cortically localised global motion mechanisms, then any transfer would reflect the broad, approximately low-pass spatial frequency tuning of global motion detectors. There would thus be most transfer when both training and test stimuli contained low spatial frequency components, and a moderate improvement for those trained or tested with broad frequency components. Since contrast discrimination is not processed in the same cortical location as global motion we predicted no transfer to contrast sensitivity would occur from any trained condition.

If transfer occurs as a result of backward generalisation, making use of the re-entrant (or feedback) connections as postulated by the Reverse Hierarchy Theory [33, 72], we predicted transfer to show specificity to the frequency tuning of the global motion processing areas. Given the broadband low-pass frequency tuning of the motion processing areas, such as V5 and V3A, we predicted robust to moderate improvement where stimuli contain low spatial frequency components (learning to transfer from low to low; modest transfer of learning from low to broad, and broad to low and broad). In addition, we predicted frequency-specific improvement in contrast sensitivity with the broad trained group showing the most transfer, and improvements reflecting the spatial frequency tuning of the training stimuli.

### Comment on the predictions

There is the possibility that transfer relies on forward connections from local to global motion mechanisms. However, given the tight frequency tuning at the local level and the well evidenced, highly specific characteristic of area V1, we would not make a prediction of transfer across frequencies occurring as a result of low level feed-forward mechanisms. However if learning did rely on the connections from local or global levels in a hierarchical fashion, we would expect frequency specific improvement on the matching spatial frequency motion condition (low to low, high to high, and broad to low and high). However, any transfer at the level of global motion, would depend on the global level mechanism so would again reflect the tuning properties of the global detectors.

### Pre- and Post-Assessment findings

Our analysis of the pre- and post-assessment data found most support for transfer as a result of backwards generalisation, however not all our predictions were supported.

Firstly, while not an explicit prediction, we had expected that the groups trained on their specific frequency conditions would naturally show improved performance for their trained frequency. However, when analysing the pre- and post-training motion results only the group trained using low spatial frequency Gabors displayed a significant learning, i.e. ‘transfer to the trained task’. Notably, the broad trained group performed worse than they did at pre-assessment stage, and there was no change for the high trained group. Furthermore the only evidence of transfer to untrained conditions, was from the low frequency trained group on the broad test condition. Pre- and post-training assessment for contrast sensitivity found that there was a significant improvement exclusively for the low trained group on the low spatial frequency contrast condition. No further improvement was found for any other trained or tested frequency. Finally, the orientation discrimination control task showed no improvement from any trained frequency.

### Evaluating the transfer from global motion

This study explored the effects of training on global motion and its transfer to other spatial frequencies and tasks. When assessing the post-training transfer to trained and untrained global motion frequencies, the *low spatial frequency trained group* was the only group to ‘transfer to trained task’ and transfer to another condition. Although the 75% threshold was worse for the broad test after training, the asymptotic performance was significantly better. Interestingly, for the high spatial frequency test, there was a significant improvement to the threshold although the slope was significantly shallower. This suggests that the sensitivity for lower coherence levels increased. Performance at the highest levels reached an asymptotic performance close to 1 at pre-assessment and remained unchanged at the post-assessment stage. Finally the shallower slope suggests a reduction in correct responses as a function of stimulus intensity.

The *high spatial frequency trained group* showed no improvement and no transfer to any other spatial frequency.

The most surprising results were those obtained from the *broad frequency trained group*. Asymptotic performance was significantly worse at the post test stage for their own trained frequency and the high frequency test, with no significant change in the low frequency test. There was no improvement in the slopes for any condition, and a small but significant improvement was found in 75% threshold for the high frequency test. Viewing individual performance revealed that the reduction in asymptotic performance may be accounted for by an outlier, however the outlier was unlikely to account for the overall absence of improvement in performance across all levels and measures, as there was no clear improvement evident for the other observers.

### Comparisons to Levi et al

Since Levi et al. (2015) reported improved contrast detection for low frequency drifting targets, we had questioned if the improvement was specific to the temporal and spatial features of the training stimuli. Half the cells in layer 4C*α* of V1, where improvement in contrast has been argued to take effect [27], are tuned for direction of motion [142]. It might be expected therefore that improvement in contrast sensitivity would be limited to moving stimuli.

Levi et al. (2015) evaluated three groups. The first trained on static, cardinally oriented Gabors, and made a horizontal or vertical direction judgement. The second trained on high intensity broadband frequency random dot coherence task, also making a direction judgement. The final group was untrained but undertook pre- and post-measures. We focus directly on the findings from the second training group which formed the basis for our initial enquiry.

#### Training

Levi et al. trained observers using a 3 up 1 down staircase, and provided auditory feedback on every trial. After 10 days training (300 trials daily), they reported an improvement in global motion perception. There was no specific pre- or post-measure for global motion integration (without trial-by-trial feedback). Improvement as a result of training was assessed by calculating the difference between performance on the first and last days of training. Our training was delivered using MOCS over 5 days with 420 trials (across 7 defined levels). To assess improvement across the training days we used a mixed-effects regression that includes all the responses over the 5 days [119]. Our training results are fully consistent with the findings of Levi et al.; training on a global motion integration task improves when training is paired with trial-by-trial feedback.

#### Pre and post-measures

As previously stated Levi et al., did not undertake pre- and post-measures for the trained task. Thus while our post-assessment for the broadband trained group showed that once feedback was removed the task was performed at prior levels, we are unable to make a direct comparison between their study and ours on this finding. Levi et al., tested four measures of transfer to contrast sensitivity; (i) orientation discrimination of static, horizontal or vertical non-flickering Gabors at varying spatial frequency (ii) orientation discrimination of moving horizontal or vertical gratings (temporal frequency of 10 hz) of varying spatial frequency (iii) direction discrimination (left/right) of moving gratings (temporal frequency of 10 hz) of varying spatial frequency and finally direction discrimination (left/right) of moving gratings (varying temporal frequency) and a spatial frequency held constant at 2 cycles per degree (medium frequency).

For (i) static ordinal Gabors, the group showed no improvement at any spatial frequency and the pre- and post-measures were no different from the untrained group. Only the moving gratings displayed an improvement in contrast sensitivity and these were predominantly found for low spatial frequency gratings.

The result from (i) is the most similar to our static contrast detection condition. Like Levi et al., our group trained on broadband global motion stimuli displayed no improvement for any tested spatial frequency for static contrast detection.

Since Levi et al. (2015) [75] reported the biggest improvement for low spatial frequency stimuli exclusively in moving targets, this may suggest improvement was specific to the temporal and spatial features of the training stimuli. In contrast, our results suggest that when trained on low frequency global motion, robust learning and transfer occurred for stimuli that contained low frequency elements, including to a static contrast sensitivity task. Both of these findings are consistent with the low-pass frequency tuning of V5.

### What our findings suggest for the mechanisms of perceptual learning

#### Specificity

We predicted that should global motion training improve contrast detection, given the frequency tuning of the early visual areas, we would expect specificity to the spatial frequency of training. Improvement was restricted to the low frequency contrast condition for the low trained group. The specificity of transfer to low frequencies is consistent with the low-pass frequency tuning of global motion areas such as V5 and V3/V3A influencing processing at V1 through feedback loops [109]. These results suggest however that there is some specificity for low frequency elements. This remains compatible with the view that global motion detectors pool via low frequency information. This may suggest that the re-entrant connections to V1, after training on global motion, only update low frequency channels.

#### Transfer

We predicted that if transfer occurred as a result of backward generalisation, then improvement would reflect the frequency tuning of the global motion detectors. The group trained on high spatial frequencies showed no improvement and no transfer. Since the representation of high spatial frequency content is attenuated in V5 [104], this may explain the lack of improvement in any post-assessment from the high or broad frequency trained groups. This may suggest any stimuli containing a high frequency element are attenuated for the purpose of the backward projections to V1 from global motion processing areas.

The broad trained group performed worse overall for almost all post-assessments. For the low spatial frequency contrast detection, there was evidence of a significantly steeper slope, however (a change in the rate of increase in proportion of correct responses with contrast), however this did not lead to a robust improvement, and the average detection threshold was worse.

Transfer was only found for the low frequency trained group. This group showed improvement in low spatial frequency contrast detection, a significant increase in asymptotic performance for the broad frequency global motion test, and also improved on their own task. This may suggest that after training on global motion only the low frequency channels are updated. The low frequency trained group will have experienced a high level of correlation in the activity in frequency channels between the feedforward and feedback connections. This joint activity may be important for perceptual learning, and is consistent with the proposal that there is an iterative interaction between the global motion processing areas and V1, since feedback from higher to lower areas engage the connections that mirror the features of the stimulus [65].

### What our findings suggest for the models of perceptual learning

The two main theories of visual perceptual learning suggest that learning either occurs through physiological changes to sensory neurons [18, 19, 31], or as a reweighting [22] of the decision units. Our findings are not able to distinguish between learning as a result of a physiological change or a higher-level computational reweighting. However neither model is able to account for the breadth of learning (and non-learning) and transfer within our results including the general absence of improvement and the occasional decline in performance at the post-assessment stage. The only difference between these components of the experiment was that trial-by-trial feedback was present for the training phase and absent for the testing phase. Our pilot study found that, for our methods, learning occurs with feedback, but not without. Additionally, Herzog and Fahle (1997) [113] suggested that once feedback is removed, performance plateaus around the level last obtained with feedback, so our expectation was that observers would maintain the improvement achieved for their trained frequency. Neither model alone accounts for the ambiguous performance at the post-assessment stage or the decline in performance by the broad trained group. On the other hand, the specificity to low spatial frequencies is consistent with both models, given the frequency tuning of the global motion detectors.

While the two positions are pitched as competing theories, they may not be mutually exclusive. In an attempt to combine the two models, Solgi and Weng (2013) [143] model learning as a two-way process (descending and ascending). The model attributes training effects to the reweighting of connections between early and higher sensory areas, but crucially also as a result of increased neuronal activity representing the trained feature [143]. For transfer to occur, Solgi and Weng (2013) [143] argue there is an important role for the re-entrant connections from higher levels. Similarly, Bejjanki et al. (2011) [144] propose a model that is computationally similar to the reweighting models, however it illustrates how changing the population codes in the early sensory areas can create similar changes in response to those made by high-level re-weighting models. Bejjanki et al. (2011) [144] argue that their model captures the characteristics for perceptual learning for both behavioural and physiological changes. Sensory inputs improve the decision weights in the feed-forward connections, and improved probabilistic inference in early visual areas as result of the increased neural activity in the feedback network. This is compatible with our findings that suggest there the high level of consistency experienced for the group trained and tested on the low frequency conditions. Ultimately, a model of perceptual learning needs to account for multiple cognitive influences. Maniglia and Seitz (2018) [145] argue that perceptual learning is not a detached or isolated process, instead multiple mechanisms react and interact to produce learning.

### Afterthoughts, Speculation and Unresolved Points

We offer some speculative ideas that may help account for the general absence of improvement at the post-assessment stage. There is emerging evidence that interleaving random stimuli may disrupt perceptual learning [146]. Yu et al. (2004) [56] found that when interleaving random contrasted Gabor stimuli, such as when using MOCS, learning did not occur, but did when only one contrast was presented at a time. Yu et al. (2004) [56] termed the interleaving of contrast stimuli as “contrast roving”, and suggest that the effect of non-learning as a result of contrast roving implies that a temporally organised pattern of stimulus presentation may be required for perceptual learning to occur. Additionally, Kuai et al. (2005) [58] found that temporal patterning was evident for motion direction discrimination. Observers were either trained using randomly varied trials or with a fixed temporal pattern of exposure (changing in a sequential clockwise direction). The first group showed no improvement in discrimination, however the second group with the fixed temporal exposure did. Cong and Zhang (2014) [147] showed that a semantic tag associated with each stimulus presentation enabled significant learning. Sequential tags such as A,B,C,D and 1,2,3,4 which contain semantic and sequential identity information both resulted in significant learning. Cong and Zhang (2014) [147] suggest that tagging and temporal sequencing may help direct the system to switch attention to the neuronal set responsible to the specific stimulus. These are consistent with the experiments conducted by Seitz et al. (2006) [136], which as previously discussed found no learning without feedback using MOCS.

In addition, there is also evidence that interleaving easy and difficult trials during perceptual training may only lead to a temporary improvement, that (a) does not persist over time and (b) disappears when easy trials are removed [148, 149]. In a difficult task without feedback learning does not occur, however it does with feedback [114]. Removing the feedback may be equivalent to removing the easy trials, and therefore reducing confidence, which is known to be an important factor in learning and perception [71]. Lin and Dosher (2017) [150] presented observers with a single block of mixed easy and difficult trials which led to rapid improvement within the block. However, after removing the easy trials and presenting only the difficult trials performance slowly returned to prior levels. This is consistent with findings that internal reinforcement could act as the teaching signal when confidence is high [34, 135].

In our study, improvement in the trained task was sustained only while feedback was present. At the post-assessment stage, the combination of a difficult task, the interleaved coherence levels and the general low decision confidence in the absence of feedback may have inhibited ‘transfer to the trained task’ and, like Lin and Dosher (2017), performance returned to baseline. It is thus a possibility that the amount of learning for the broad and high spatial frequency trained groups was insufficient to overcome the reduction in performance when feedback was removed.

An exception to this was the improvement in performance and ‘transfer to the trained task’, for the low frequency trained group. Since learning and transfer to trained task were evident for the low spatial frequency trained group, this may suggest an interaction between the amount of learning, and the benefit of feedback. Furthermore, the improvement by the low frequency group over the five training days shows the biggest move from baseline performance, and thus potentially benefited from more learning, which was robust, persistent and thus incurred less disruption by the absence of feedback.

## Conclusions

Our results support the early ideas from Reverse Hierarchy Theory that the feedback connections from higher levels back to early visual processing areas may be involved in facilitating learning that transfers [33, 72], and are consistent with models that suggest that the re-entrant connections play a vital role in learning and transfer [143, 144].

Our first experiment suggested that for perceptual learning of an equivalent noise global motion task, presented using MOCS, feedback was a necessity. However after training with feedback, there was little evidence of a robust improvement when feedback was removed at the post-assessment stage. The only condition that showed unambiguous learning and transfer was the low spatial frequency trained group. Our findings are in line with current research which finds global motion detectors pool for low spatial frequencies.

Our main experiment found robust learning and transfer for the group trained with low spatial frequency specific global motion gabors. This has provided evidence of frequency specific transfer from a higher-level global motion task to a low-level static untrained local task. The frequency tuning of these results is consistent with perceptual learning which depends on the global stage of motion processing.

Finally, the frequency tuning of these results suggest that the mechanism of transfer of global motion is dependent on backward generalisation, and specific to the frequency tuning of the global motion detectors. Transfer exhibited a high specificity towards low spatial frequencies, consistent with the attenuation found in the processing of global motion.

Our study supports the idea that the perception of global motion shows a bias for low spatial frequencies and suggests the backward projections and the re-entrant connections from global motion processing areas to V1 play a crucial role in transfer to trained and untrained tasks.

## Acknowledgments

This research was funded by a University of Essex Doctoral studentship, and grants from ESSEXLab and PsyPAG to J.A.

